# SIRT3 deficiency decreases oxidative-metabolism capacity but increases lifespan under caloric restriction

**DOI:** 10.1101/2022.05.09.491205

**Authors:** Rashpal S Dhillon, Yiming (Amy) Qin, Paul R van Ginkel, Vivian X Fu, James M Vann, Alexis J Lawton, Cara L Green, Fúlvia B Manchado-Gobatto, Claudio A Gobatto, Dudley W Lamming, Tomas A Prolla, John M Denu

**Affiliations:** Department of Biomolecular Chemistry, University of Wisconsin-Madison, Madison, WI, USA; Wisconsin Institute for Discovery, University of Wisconsin-Madison, Madison, WI, USA; Interdisciplinary Graduate Program in Nutritional Sciences, University of Wisconsin-Madison, Madison, WI, USA; Department of Genetics and Medical Genetics, University of Wisconsin-Madison, Madison, WI, USA; Department of Medicine, SMPH, University of Wisconsin-Madison, Madison, WI, USA; William S. Middleton Memorial Veterans Hospital, Madison, WI, USA; School of Applied Sciences, Laboratory of Applied Sport Physiology, University of Campinas, Limeira, Brazil

**Keywords:** calorie restriction, lifespan, mitochondrial acetylation, mitochondrial respiration, fatty acid oxidation, aerobic fitness, fuel switching, pseudo-fasting

## Abstract

Mitochondrial NAD^+^-dependent protein deacetylase Sirtuin3 (SIRT3) has been proposed to mediate calorie restriction (CR)-dependent metabolic regulation and lifespan extension. Here, we investigated the role of SIRT3 in CR-mediated longevity, mitochondrial function, and aerobic fitness. We report that SIRT3 is required for whole-body aerobic capacity but is dispensable for CR-dependent lifespan extension. Under CR, loss of SIRT3 (*Sirt3^-/-^*) yielded a longer overall and maximum lifespan as compared to *Sirt3^+/+^* mice. This unexpected lifespan extension was associated with altered mitochondrial protein acetylation in oxidative metabolic pathways, reduced mitochondrial respiration, and reduced aerobic exercise capacity. Also, *Sirt3^-/-^* CR mice exhibit lower physical activity and favor fatty acid oxidation during the postprandial period, leading to a pseudo-fasting condition that extends the fasting period. This study shows uncoupling of lifespan and healthspan parameters (aerobic fitness and spontaneous activity), and provides new insights into SIRT3 function in CR adaptation, fuel utilization, and aging.

## Introduction

Incidence of non-communicable diseases, such as cardiovascular disease, neurodegeneration, and cancer increase significantly with age (Balasubramanian, Howell and Anderson, 2017). CR robustly delays age-related diseases and extends both health and lifespan in diverse species (Fontana and Partridge, 2015; Mattison *et al*., 2017). Mitochondria, the center of cellular oxidative metabolism, are vital to cellular health and mitochondrial dysfunction has been associated with accelerated aging (Srivastava, 2017; Jang *et al*., 2018). One of the hallmarks of CR is the preservation of mitochondrial function through reducing oxidative stress, enhancing fuel utilization, and maintaining mitochondrial dynamics and integrity (Merry, 2004; Bruss *et al*., 2010; Lanza *et al*., 2012; Jang *et al*., 2018). Previous studies have proposed that SIRT3-dependent deacetylation may play a major role in modulating mitochondria under CR (Qiu *et al*., 2010; Someya *et al*., 2010; Hallows *et al*., 2011).

Reversible N^ε^-lysine acetylation is a prominent post-translational modification enriched in mitochondria, with more than 60% of all mitochondrial proteins having at least one acetylated lysine site identified in proteomic studies (Liu *et al*., 2014; Baeza, Smallegan and Denu, 2016). Among well-characterized examples, mitochondrial acetylation mostly inhibits enzymatic activity, slows down metabolic pathways, and leads to mitochondrial dysfunction (Hirschey *et al*., 2010; Still *et al*., 2013; Vassilopoulos *et al*., 2014) which has been correlated with diabetes, heart failure, and age-related cellular impairment (Fukushima and Lopaschuk, 2016; Horton *et al*., 2016; Ansari *et al*., 2017). SIRT3 is the predominant NAD^+^-dependent deacetylase in mitochondria, whose deficiency leads to significant hyperacetylation in mitochondria of various tissues (Dittenhafer-Reed *et al*., 2015). SIRT3 level is diet sensitive, and its expression increases under fasting and CR (Palacios *et al*., 2009; Schwer *et al*., 2009). We have reported SIRT3-dependent deacetylation of mitochondrial proteins in CR mice, and demonstrated Sirt3 is essential for the prevention of age-related hearing loss in mice fed a CR diet (Someya *et al*., 2010). These observations have fueled the speculation that SIRT3- dependent control of acetylation may be crucial for CR-induced modulation of aging and lifespan extension. Nonetheless, direct evaluation of the contribution by Sirt3 to CR-dependent lifespan extension and mitochondrial performance during aging is lacking.

Here, using male Sirt3*^+/+^* (WT) and *Sirt3^-/-^* mice, both in a C57BL/6J/*Nnt^+/+^* genetic background, treated with either a control diet (CD) or CR diet, we found that *1*) SIRT3 is essential for whole-body aerobic fitness, but is not required to mediate CR-dependent longevity, *2*) loss of SIRT3 reduces the net OXPHOS respiration when glucose-derived metabolites are utilized, and *3*) SIRT3 ablation promotes fuel switching from glucose to fatty acids during the postprandial state. We also show that SIRT3 is required for CR-induced increases in spontaneous physical activity, effectively lowering whole body energy expenditure of *Sirt3^-/-^* mice under CR. Collectively, these data reveal that SIRT3 deficiency reduces the capacity to oxidize carbohydrates under CR and that this altered mitochondrial metabolism favoring fatty acid metabolism is associated with an increase in lifespan. We introduce the idea of ‘pseudo fasting’ and its possible link to increased lifespan, given the emerging data that fasting duration in a CR regime may be a determinant of longevity.

## RESULTS

### SIRT3 ablation under CR extends lifespan and has minimal impact on mitochondrial integrity and ROS detoxification

To investigate the role of SIRT3 in CR-dependent metabolism during aging, we established whole-body *Sirt3^-/-^* mouse models in the *Nnt^+/+^* background by initially crossing C57BL/J (*Nnt^-/-^) Sirt3^-/-^* mice with C57BL/6N (*Nnt^+/+^*) mice. This is noteworthy in that many prior aging and SIRT3 studies employed C57BL/6J mice that lack functional mitochondrial Nicotinamide Nucleotide Transhydrogenase (NNT). With increasing evidence demonstrating the importance of NNT in redox control and metabolism (Freeman *et al*., 2006; Ronchi *et al*., 2016; Leibel R, Rosenbaum M, 2020), we generated and used C57BL/6J/*Nnt^+/+^* mice in the current study. From 2 months of age until death, male WT and *Sirt3^-/-^* mice were maintained on a CD or CR diet (25% less calorie intake than control diet fed mice) (Fig.1A). The use of ∼15% less calories compared to ad libitum in CD fed mice was designed to reduce obesity and related metabolic complications (Pugh, Klopp and Weindruch, 1999).

**Fig 1.**
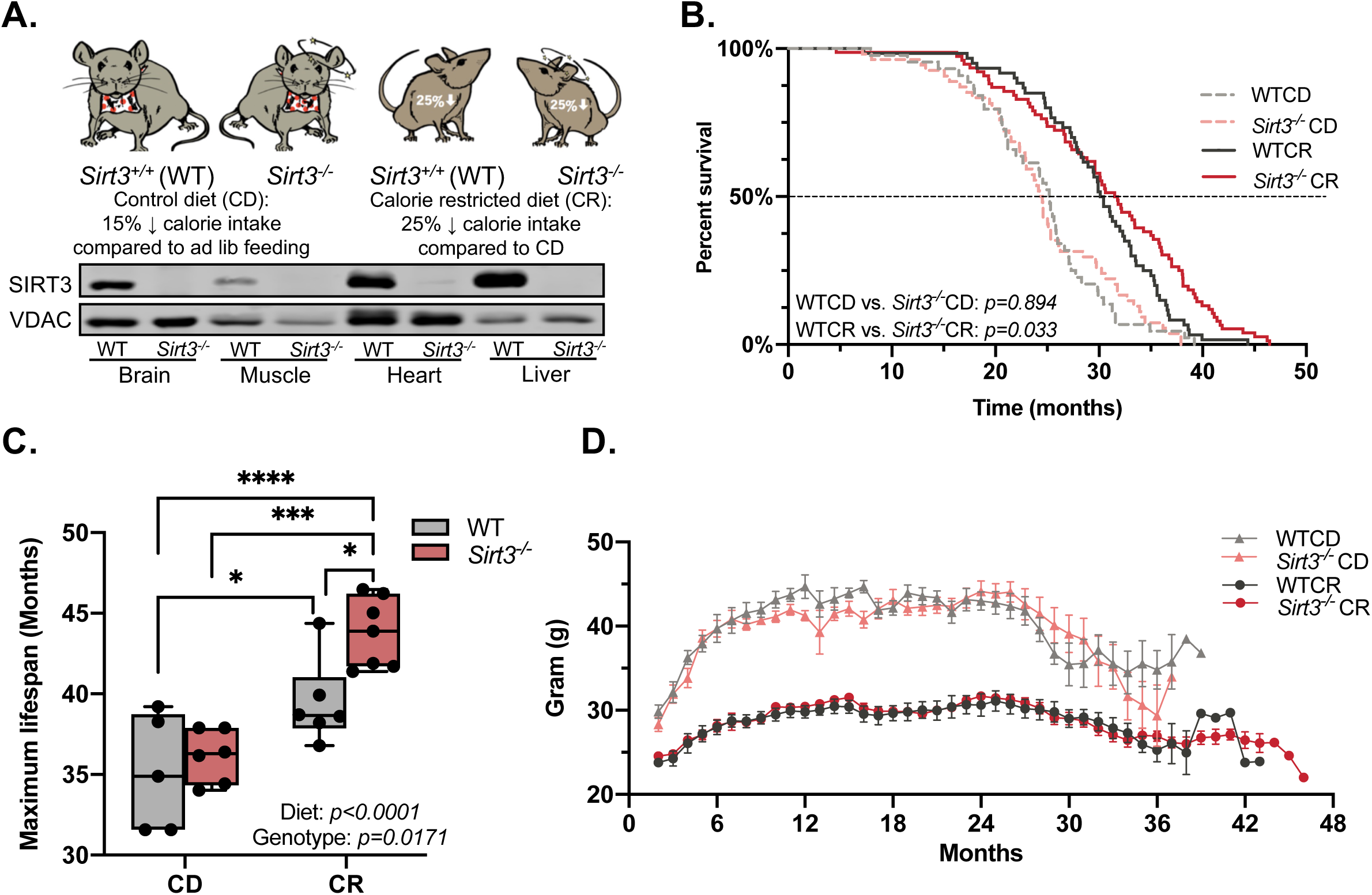
**Sirt3 ablation under CR extends lifespan.** A) Mouse models in the current study. B) Kaplan-Meier survival curves for WTCD (n=44), WTCR (n=60), *Sirt3^-/-^*CD (n=54), and *Sirt3^-/-^* CR mice (n=76). The dashed line (50% survival probability) indicates the median lifespan. Log-rank test was used to calculate statistical significance of lifespan among treatment groups. See Table S1A for detailed statistics. C) Maximum lifespan is represented by the average lifespan of the top 10% longest lived mice for each group. Data were analyzed by two-way ANOVA followed by multiple comparisons test. *p* value reported for each comparison is corrected by Tukey’s test. Results are plotted as mean with min. to max. range. *: *p*≤0.05; **: *p≤*0.01; ***: *p≤*0.001; ****: *p*≤0.0001. We also confirmed the statistical significance of maximum lifespan between WTCR and *Sirt3^-/-^*CR using Boschloo’s Test (Wang-Allison). See Table S1B for details. D) Lifetime body weight for mice used in lifespan study shown in Fig.1B.

Kaplan-Meier survival curves (Fig.1B, Table S1A) revealed that as expected, CR significantly extended the average, median (50% survival), and maximum lifespan (Fig.1C, calculated as the average lifespan of the top 10% longest lived mice) of WT mice. *Sirt3^-/-^*mice exhibited a similar average, median lifespan as their diet-matched counterparts under both CD and CR, but unexpectedly *Sirt3^-/-^* CR mice showed an overall increase in lifespan as compared to WTCR mice (*p*=0.033). *Sirt3^-/-^* CR mice displayed a 11% increase in maximum lifespan relative to WTCR (Fig.1C. Confirmed with Boschloo’s Test, *p*=0.0405 at 90% percentile for WTCR vs. *Sirt3^-/-^* CR, Table S1B). These results show that SIRT3 is not required for the normal life extension benefit of CR and that SIRT3 ablation further extends CR-mediated longevity. Interestingly, the *Sirt3^-/-^*-dependent overall lifespan extension benefit was not noticeable until a later age, as the survival curves of WTCR and *Sirt3^-/-^*CR mice diverge after the median lifespan of these groups (Fig.1B) and a significant change in 75% percentile in WTCR vs. *Sirt3^-/-^* CR was observed (Table S1B). Lifetime body weight (Fig.1D) and body composition at 25 months of age (Fig.S1A-C) showed a diet- but not a SIRT3-dependence. Together, these observations revealed an unexpected boost of maximum lifespan in *Sirt3^-/-^* mice under CR.

Several reports have highlighted the ability of SIRT3 to preserve mitochondrial integrity and suppress oxidative stress through protein deacetylation (Kim *et al*., 2010; Qiu *et al*., 2010; Bause and Haigis, 2013; Ono *et al*., 2019) including work demonstrating that CR prevents age-related hearing loss through SIRT3-dependent mechanism involving IDH2 activation (Someya *et al*., 2010; Yu, Dittenhafer-Reed and Denu, 2012). Recently, Benigni et al. (2019) observed that shortened lifespan in standard chow fed *Sirt3^-/-^* mice is associated with altered cardiac mitochondrial morphology. Given the unexpected longevity in *Sirt3^-/-^* CR mice, we investigated whether mitochondrial integrity as examined by transmission electron microscopy (TEM) were affected at 25 months of age. No morphological differences in gastrocnemius and heart mitochondria were observed between genotypes or diets (Fig. S1D). Similarly, citrate synthase activity, a surrogate marker of mitochondrial density, and its protein expression in the brain, gastrocnemius, heart and liver were comparable between groups (Fig.S1E-F). Moreover, there was no significant link between SIRT3 expression (Fig.S1G) and mitochondrial DNA content (mtDNA:nDNA, Fig.S1H). NADPH/NADP^+^ and GSH/GSSG ratios were comparable between *Sirt3^-/-^* and WT animals under both diets (Fig. S1I-J) in the liver and heart. In sum, these data indicate that loss of SIRT3 has no significant impact on mitochondrial integrity and markers of ROS detoxification in aged C57BL/6J/*Nnt^+/+^* mice.

### SIRT3 opposes hyperacetylation of its targets under CR during aging

Consistent with the role of SIRT3 as a mitochondrial deacetylase, immunoblotting of total mitochondrial protein acetylation in liver from 25-month-old mice displayed an increase with SIRT3 ablation and a combined effect of CR and loss of SIRT3 (Fig. 2A). To identify the lysine sites and proteins showing altered acetylation as a function of age, genotype and diet, we leveraged our recently developed quantitative mass spectrometry workflow (Baeza *et al*., 2020) (Fig. S2A). Site-specific acetylation stoichiometry from enriched liver mitochondrial fractions was determined in 5- and 25-month-old WTCD, WTCR, *Sirt3^-/-^*CD and *Sirt3^-/-^*CR mice. From 1,854 unique mitochondrial acetyl-lysine sites identified (≤1% false discovery rate), 941 sites on 329 proteins (MitoCarta 2.0 (Calvo, Clauser and Mootha, 2016), Table S2) were quantified in all 8 treatment groups, with acetylation stoichiometry ranging from less than 1% to 99% and a median stoichiometry of ∼8.3% amongst quantified sites.

**Fig 2.**
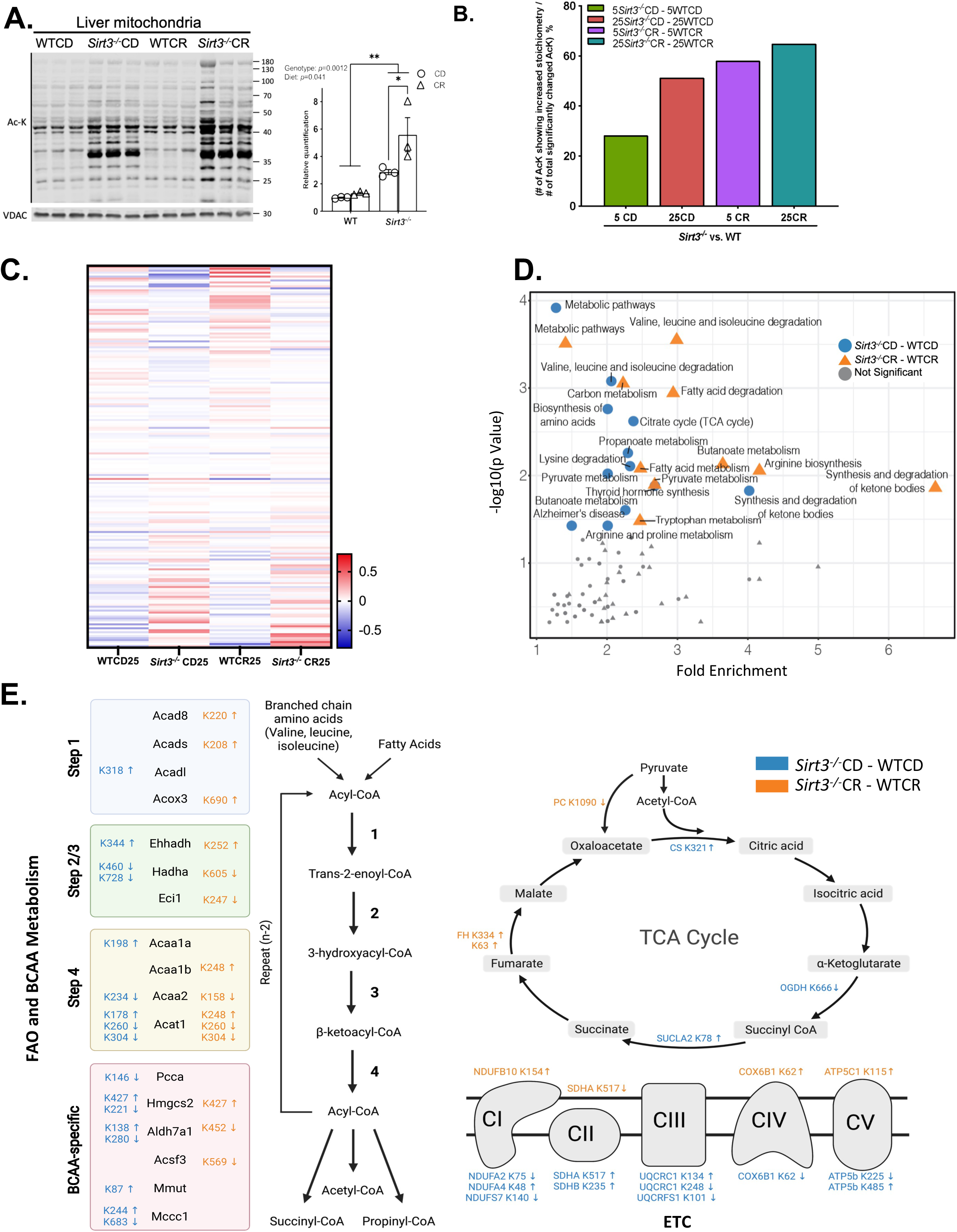
A) Immunoblot of pan-acetylation, normalized to VDAC, for liver mitochondrial enrichment from 25-month-old treatment groups. Data were analyzed by two-way ANOVA followed by multiple comparisons test. *p* value reported for each comparison is corrected by Tukey’s test. Results are plotted as mean ± SEM. *: p≤0.05. B) Percentage of significantly changed acetyl-lysine residues that show increased stoichiometry due to *Sirt3^-/-^*, calculated by (the number of acetyl-lysine sites showing increased stoichiometry) / (the number of significantly changed acetyl-lysine sites, p<0.05) x100%. C) Heat map of significantly changed lysine sites (p<0.1) that are response to loss of SIRT3. Plotted sites are significantly changed (p<0.1) in either *Sirt3^-/-^*CD vs. WTCD or *Sirt3^-/-^*CR vs. WTCR comparison. Values are colored based on relative acetylation stoichiometry, normalized to the median value of each sites in all four groups, scaling ranging from -0.8 to 0.8 (x100%). D) Functional cluster analysis of KEGG pathways (DAVID 6.8). Significantly enriched (-log10(p value) >1.5) pathways are indicated, with *Sirt3^-/-^*CD vs. WTCD in orange and *Sirt3^-/-^*CR vs. WTCR in blue. E) Acetylation sites in FAO and BACC metabolism, TCA cycle, and ETC that displayed larger than 5% stoichiometry (p<0.1) for *Sirt3^-/-^*CD vs. WTCD (orange colored) and *Sirt3^-/-^*CR vs. WTCR comparison (blue colored).

The effects of diet, age and genotype were parsed out (Figs.S2B-C) to reveal the relative number of changed acetylation sites (density) as a function of change in stoichiometry for those conditions. An age-mediated increase in acetylation stoichiometry in mice fed a CR diet was evident. Both young WT and *Sirt3^-/-^*CR mice showed less acetylation alteration and accumulation, suggesting that aged mice are more prone to CR-induced hyperacetylation (Fig.S2C). Additionally, the stoichiometry disparities between aged and young mice were more notable when the mice are *Sirt3^-/-^* and CR fed, indicating that both CR and *Sirt3^-/-^* amplify the age-dependent acetylation alteration. Lastly, SIRT3-dependent acetylation changes are most significant in aged, CR-treated animals (Fig. S2E), with the 25-month *Sirt3^-/-^*CR vs. 25-month WTCR comparison showing the largest fraction of increased acetylation (Fig. 2B) amongst the various groups. In sum, these results indicate that CR, and loss of SIRT3 individually and cooperatively contribute to the global increase in acetylation at the old age.

To reveal specific site-level acetylation changes across the various conditions, we plotted the relative stoichiometry of significantly changed sites due to loss of SIRT3 in WTCD vs. *Sirt3^-/-^*CD or WTCR vs. *Sirt3^-/-^*CR comparisons at 25 months (Fig. 2C). As noted in the major acetylation trends displayed in Fig. S2C-E, many sites showed increased acetylation in *Sirt3^-/-^* mice and an additive effect in *Sirt3^-/-^*CR. These sites likely reflect direct SIRT3 targets that become hyper-acetylated under CR. Most surprisingly, we identified sites which were hypoacetylated in *Sirt3^-/-^* mice versus WT counterparts under both CD and CR (*Sirt3^-/-^*CD vs. WTCD: 65 sites, *Sirt3^-/-^*CR vs. WTCR: 31 sites), and sites that were hypoacetylated in *Sirt3^-/-^*CR vs. WTCR but not *Sirt3^-/-^*CD vs. WTCD (11 sites) (Table S2). These sites show opposite behaviors to those expected, but yet their changes in acetylation follow a SIRT3-dependent manner. This is especially interesting for sites that are hypoacetylated only in CR when Sirt3 is absent. These observations reveal a set of acetylation sites that are dependent on CR and SIRT3, but would not be direct substrate targets of SIRT3. The acetyl-proteomics indicate major changes to the acetylation status of mitochondrial proteins dependent on age, genotype and diet, and that the unique set of expected and unexpected changes under both CR and loss of SIRT3 may highlight the molecular pathways in *Sirt3^-/-^*CR mice associated with increased lifespan.

To identify the mitochondrial processes that are likely perturbed due to *Sirt3^-/-^* and CR at an old age, we performed functional cluster analysis of KEGG pathways (Fig.2D, DAVID 6.8) (Dennis *et al*., 2003). We compared the KEGG pathway enrichment analysis and found many similarities between the enriched pathways in the CD fed *Sirt3^-/-^*, WT mice and the CR fed *Sirt3^-/-^*, WT mice. In both comparisons (Fig. 2D), significant enrichment of major metabolic pathways was apparent, including TCA cycle, valine, isoleucine and leucine degradation, and fatty acid degradation. In the most affected pathways (Fig. 2E), 52 acetylation sites (>5% stoichiometry change, p<0.1) were identified in fatty acid oxidation, TCA cycle, electron transport chain (ETC), and branched chain amino acids (BCAA) metabolism. Amongst these sites, 23 sites were previously noted in Hebert *et al*., (2013) and 29 sites are newly identified. In addition to increased acetylated sites due to loss of SIRT3, we observed 44% of the sites displaying lower acetylation stoichiometry in *Sirt3^-/-^*CD and CR mice relative to their WT counterparts, again suggesting that alteration in acetylation occurred in unexpected combinations of sites that decreased or increased in acetylation. Regardless of direction change in acetylation, the cluster analysis suggests a functional alteration in oxidative metabolism is likely associated with the extended lifespan observed in *Sirt3^-/-^*CR mice (Fig. 2D-E).

### Loss of SIRT3 limits aerobic fitness due to reduced mitochondrial respiration capacity

To understand how altered acetylation of pathways identified in the acetyl-proteomics might affect mitochondrial oxidative metabolism in aged *Sirt3^-/-^*CR mice, we first assessed whole-animal bioenergetics through aerobic exercise capacity. At 25 months of age, WTCD, *Sirt3^-/-^*CD, WTCR, *Sirt3^-/-^*CR mice were subjected to 4 sessions of treadmill exercise at different velocities until exhaustion, and the critical velocity, a measure of aerobic exercise capacity, was obtained (Scariot *et al*., 2019). Both *Sirt3^-/-^* groups performed poorly, displaying critical velocity for *Sirt3^-/-^*CD and *Sirt3^-/-^*CR mice that were 24% and 27% slower than that of their diet matched control animals (Fig.3A).

**Fig 3.**
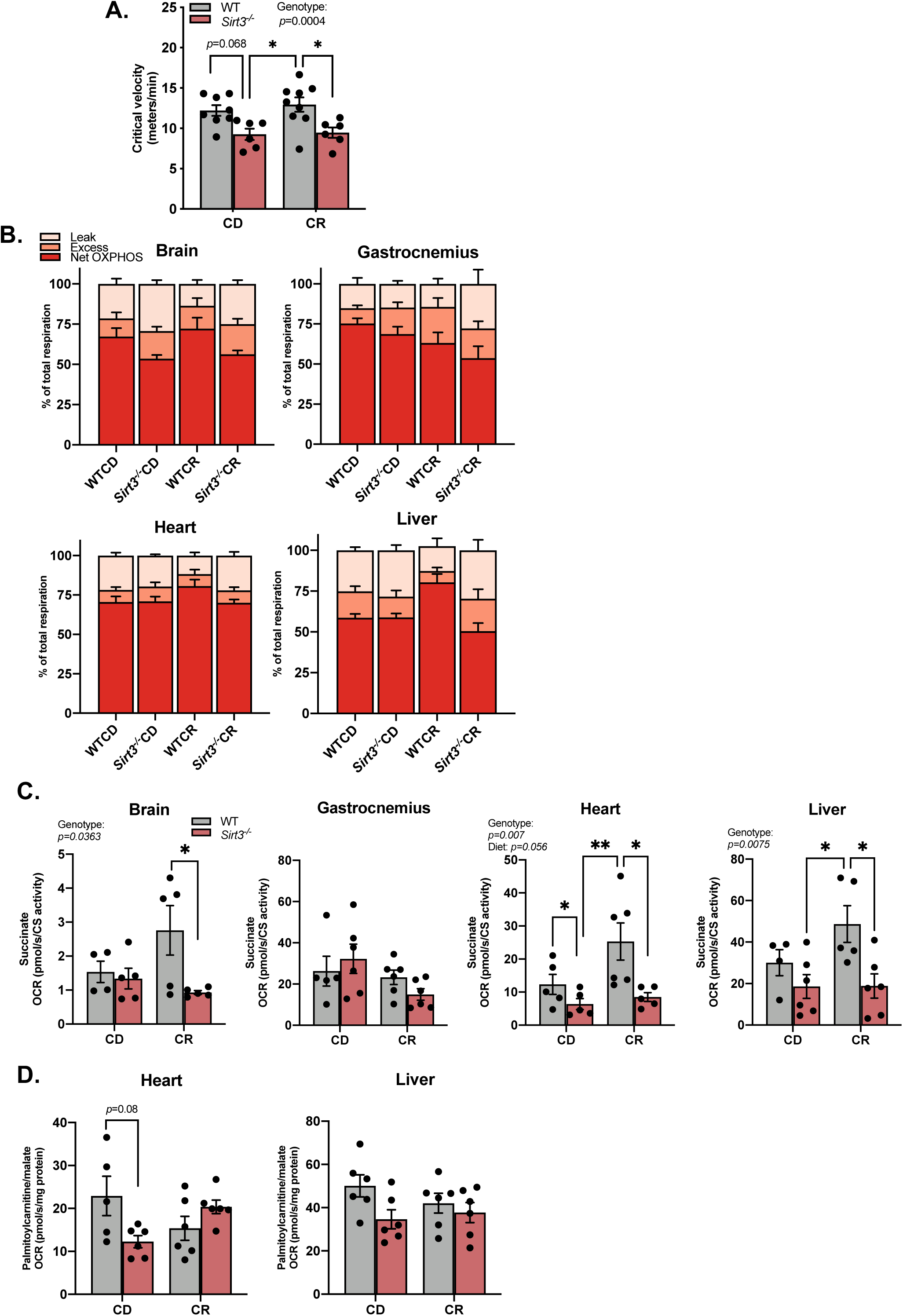
**Loss of SIRT3 limits aerobic fitness due to reduced mitochondrial respiration capacity** A) The critical velocity of 25-month-old WTCD, *Sirt3^-/-^*CD, WTCR, *Sirt3^-/-^* CR mice, n=6-9 per group. B) Percentages of leak, excess and net OXPHOS respiration to total respiration in permeabilized brain, gastrocnemius, heart and liver from 24 hours fasted 25-months-old mice, n= 5-6 per group. Each of these respiration parameters were assessed by oxygen consumption rate (OCR). Total respiration (electron transfer capacity) was assayed using pyruvate/ glutamate/ malate/ succinate as substrates upon mitochondrial uncoupler FCCP addition. Leak respiration was assayed using pyruvate/ glutamate/ malate in the absence of ADP. Coupled respiration was assayed using pyruvate/ glutamate/ malate/ succinate in the presence of ADP. Net-OXPHOS respiration was calculated by subtracting leak respiration from coupled respiration. Excess respiration is calculated by subtracting coupled respiration from electron transfer capacity. Percentages of leak, excess and OXPHOS respiration to total respiration were calculated by (OCR of each respiration parameter)/(ORC of total respiration) x100%. See Fig. S3A for OCR of each respiration parameter and Fig.S3B for statistical details. C) Succinate-dependent respiration capacity was assessed using succinate as substrate in the presence of FCCP and rotenone. D) FAO-dependent coupled respiration was assayed in liver and heart from 24 hours fasted 25-months-old mice using palmitoylcarnitine/malate as substrates in the presence of ADP. Data were analyzed by two-way ANOVA followed by multiple t-tests. *p* value reported for each comparison is corrected by Tukey’s test. Results are plotted as mean ± SEM. *: *p*≤0.05; **: *p≤*0.01. Significant (p≤0.05) diet effect, genotype effect and/or diet and genotype interaction for each experiment are indicated in figures.

Next, we investigated whether reduced aerobic fitness in *Sirt3^-/-^* mice can be explained by compromised mitochondrial respiration given that subunits of ETC complexes were one of the most prominent SIRT3 target groups and found to be inhibited under hyperacetylation induced by SIRT3 ablation (Ahn *et al*., 2008; Bao *et al*., 2010; Horton *et al*., 2016; Parodi-Rullán, Chapa-Dubocq and Javadov, 2018) (Table S2). To this end, we measured *ex-vivo* mitochondrial respiration in permeabilized brain, gastrocnemius, heart, and liver from 25-month-old treatment groups. Using high resolution respirometry with pyruvate/glutamate/malate/succinate as substrates, we found that WTCR mice exhibited the highest electron transport capacity in heart (Fig.S3A) and the highest percentage of net OXPHOS respiration relative to total respiration in heart and liver (Fig.3B, see Fig.S3B for statistics) among four treatment groups, consistent with previous reports that CR preserves respiration efficiency in aged mice (Weindruch *et al*., 1980; Lanza *et al*., 2012). In the CD group, net OXPHOS respiration (Fig.S3A) and its percentage to total respiration (Fig.3B) are largely comparable in gastrocnemius muscle, heart and liver between genotypes, with a ∼20% reduction (*p*=0.042) in percentage of net OXPHOS respiration found in brain mitochondria from *Sirt3^-/-^* relative to WT mice. In contrast, under CR, we observed pronounced reductions in net OXPHOS respiration (Fig.3B) and its percentage relative to total respiration capacity (Fig.S3A) in brain, heart and liver in *Sirt3^-/-^*CR compared to WTCR mice. These respiration phenotypes suggest that both SIRT3 and CR can play a role in preserving mitochondrial respiration capacity and bioenergetics in old age, and that reduced net OXPHOS respiration is a consistent feature of *Sirt3^-/-^*CR mice.

To pinpoint the affected respiration pathway(s), we parsed out substrate-dependent respiration and conducted enzymatic activity assays for the 25-month-old treatment groups. No significant difference in NADH-linked substrates (pyruvate/glutamate/malate) coupled respiration between genotypes nor between diets was observed (Fig.S3C). Succinate-dependent respiration capacity, however, was significantly reduced in *Sirt3^-/-^*CR liver, and heart relative to WTCR (Fig.3C). In CD treated groups, succinate-dependent respiration capacity showed minor difference in heart between WT and *Sirt3^-/-^*. Enzymatic activity assays revealed that SIRT3 and CR indeed had the more prominent impact on Complex II activity compared to that of Complex I (Fig.S3D-E). Moreover, we determined if fatty acid oxidation (FAO)-dependent respiration was limited in permeabilized liver and heart from *Sirt3^-/-^*animals using palmitoylcarnitine/malate as substrates. Interestingly, palmitoylcarnitine-dependent coupled respiration (Fig.3D) and its net OXPHOS control efficiency (Fig.S3F) was similar between WTCR and *Sirt3^-/-^*CR mice in both liver and heart. In contrast to mice on the CD diet, palmitoylcarnitine-dependent OXPHOS control efficiency (Fig. S3F) was significantly lower in *Sirt3^-/-^*mouse heart relative to WT. These results suggest that the reduced Complex II respiration in SIRT3-deficient mice provides a molecular basis for lowered whole-body aerobic exercise capacity. But notably, CR treatment allows *Sirt3^-/-^*mice to maintain FAO-dependent respiration.

### *Sirt3^-/-^* mice under CR display faster switching from glucose to fatty acid oxidation during the postprandial period

In light of the critical speed and substrate-dependent respiration phenotypes obtained from the four treatment groups, we speculated that *Sirt3^-/-^*CR mice display unique metabolic features. To this end, the four groups of 25-month-old mice were subjected to a metabolic chamber experiment that consisted of a 24-hour fasting period followed by a refeeding period with 8 hours of food provision (Fig.4A-B). Diet-matched groups consumed a similar amount of food during the 8-hour food provision period (Fig.4C). During the fasting period, the respiratory exchange ratio (RER) was not significantly different among the treatment groups, with RERs ranging between 0.7-0.8, indicating that fatty acids were the preferred energy source. Upon feeding, RER increased to ≥1 for all groups, reflecting a fuel switching to predominant use of carbohydrate as an energy source, accompanied by fatty acid synthesis before returning to FAO in the fasted state (Bruss *et al*., 2010; Mitchell *et al*., 2019). *Sirt3^-/-^*CR mice displayed smaller amplitudes of RER elevation upon feeding and a shorter duration of the RER≥1 period as compared to that of WTCR mice, whereas WTCD and *Sirt3^-/-^*CD showed no difference in these parameters. RER, fed and fast FAO (Fig.4B, D-E) suggest an increased trend in FAO in *Sirt3^-/-^*CR compared to WTCR mice.

**Fig 4.**
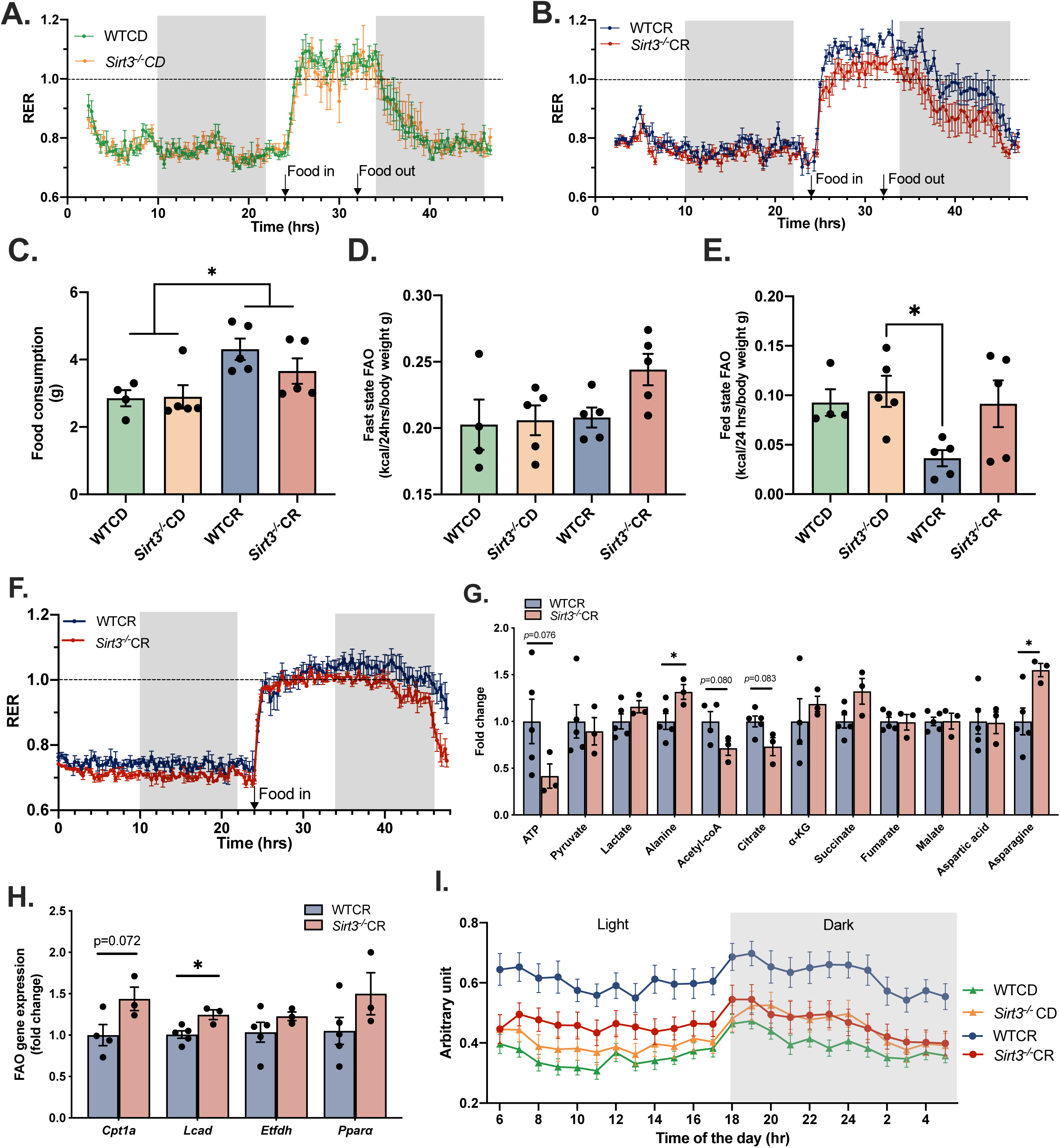
***Sirt3^-/-^* mice under CR display faster switching from glucose to fatty acid oxidation during the postprandial period** A-C) Respiratory exchange ratio (RER) and food consumption of 25-month-old WTCD, *Sirt3^-/-^*CD, WTCR, *Sirt3^-/-^* CR mice in the metabolic chamber experiments, n=5. Duration of food availability is indicated in both figures, which is the interval between food in and food out. D-E) FAO during fast and fed states for mice shown in Fig.4A-B. Data reported here are corrected to body weight. F) Respiratory exchange ratio (RER) of 25-month-old WTCR, *Sirt3^-/-^* CR mice, n= 5,3. Food was provided at the 24^th^ hour of the experiment, indicated as food in. The normal daily amount of food was provided, and no food was taken out during the experiment. G) Fold change of major TCA metabolites in 25-month-old CR-treated mice liver after a 6-hour refeeding period, n= 5 (WTCR), 3 (*Sirt3^-/-^* CR). H) Fatty acid oxidation gene expression in 25-month-old CR-treated mice liver after a 6-hour refeeding period. Gene expressions is normalized to β-2-microglobulin and relative to WTCR values, n= 5 (WTCR), 3 (*Sirt3^-/-^* CR). I) Spontaneous physical activity in arbitrary units at the light and dark phases for 25 months WTCD, *Sirt3^-/-^*CD, WTCR, *Sirt3^-/-^* CR mice over a 19-day period, n=6 per group. Each data point represents the 19-day average of spontaneous activity of all mice in the same treatment group at the indicated hour of the day. Data were analyzed by one-way ANOVA followed by multiple t-tests. P value reported for each comparison is corrected for multiple comparisons using the Bonferroni test. Results are plotted as mean ± SEM. *: *p*≤0.05; **: *p≤*0.01;

Given the every-other-day feeding protocol in the current lifespan experiment for both CD and CR animals (Pugh, Klopp and Weindruch, 1999), CR animals generally consumed their daily allotment of food within 12-16 hours without altered eating behavior observed between genotypes (See Methods for the detailed feeding protocol). To capture daily RER fluctuations and estimate the duration of the predominant carbohydrate utilization period in CR mice, we repeated the metabolic chamber experiments using a different set of CR animals with 24 hours of fasting followed by a refeeding period in which the normal daily food allotment used in the lifespan study was provided, and no food was removed during the experiment. These conditions mimic the normal food intake routine of this cohort of animals throughout their lifespan. Again, a smaller amplitude of RER in response to food intake and a shorter duration of RER ≥1 were observed in *Sirt3^-/-^*CR relative to WTCR mice (Fig.4F, S4A-B), consistent with the RER pattern observed in Fig.4A-B. These observations suggest that *Sirt3^-/-^*CR mice are able to switch to carbohydrate as the predominant fuel source upon feeding, but are less capable of maintaining carbohydrate utilization during the fed state. As a result, *Sirt3^-/-^*CR animals experience an earlier switch to FAO-dependent metabolism (fasting state) compared to WTCR. Here, we introduce the concept of “*pseudo-fasting”* to represent this faster switching to the FAO period that occurs in the fed state of *Sirt3^-/-^*CR mice.

To further investigate whether *Sirt3^-/-^*CR mice have compromised glucose oxidation during the fed state, we performed LC-MS based metabolite profiling in liver, heart and muscle of 6-hour refed mice. Compared to WTCR mice, the *Sirt3^-/-^*CR group exhibited a trend of reduced levels of ATP (*p*=0.07), acetyl-CoA (*p*=0.08) and citrate (*p*=0.08) accompanied with a significant accumulation of alanine and asparagine in liver (Fig.4G), and reduction of citrate and accumulation of asparagine (p=0.064) in heart (Fig.S4C). Reduced pyruvate and α-KG were observed in heart of *Sirt3^-/-^*CD compared to WTCD (Fig. S4E), and no difference was detected in liver (Fig. S4D). No difference in TCA metabolites was found in gastrocnemius among all groups except an increased pyruvate noted in CR groups (Fig.S4F). Next, we examined whether FAO genes were upregulated at an earlier time upon feeding, which could explain the decline of the RER in *Sirt3^-/-^*CR mice. We measured transcript levels of key FAO genes, including *Cpt1*, *Lcad*, *Etfdh*, and *Pparα* in 6-hour refed liver from 25-month-old WTCR and *Sirt3^-/-^*CR mice (Fig. 4D). After 6 hours of food provision, *Sirt3^-/-^*CR mice displayed trends of increased FAO gene expression, consistent with *Sirt3^-/-^*- and CR-dependent fuel utilization adaptation observed in whole body RER studies.

Next, we investigated how *Sirt3^-/-^*CR mice could maintain a similar body composition (Fig.S1B-D) to that of WTCR mice given the ability of *Sirt3^-/-^*CR to switch toward FA oxidation more quickly and shorter lipogenesis period (RER≥1, Talal *et al*., 2021). We examined spontaneous physical activity (SPA) as an important component of daily energy expenditure (Manini, 2010). Here, we observed that loss of SIRT3 blunted CR-mediated higher spontaneous physical activity in a 19-day recording period (Fig.4E). These results indicate that SIRT3 is required for the higher spontaneous physical activity under CR and suggest that *Sirt3^-/-^*CR mice expend significantly less energy, allowing these mice to preserve comparable fat mass to WTCR animals. The net consequence is that SIRT3 loss under CR results in a long-lived animal that is less active, has reduced exercise and oxidative capacity due to attenuated carbohydrate oxidation capacity, and preferentially utilizes fat as energy source.

## Discussion

CR is a widely studied regimen in mice that robustly extends both healthspan and lifespan (Weindruch *et al*., 1986; Zhang *et al*., 2013; Fontana and Partridge, 2015; Mitchell *et al*., 2019). Previous work revealed that CR can stimulate SIRT3 expression (Palacios *et al*., 2009; Schwer *et al*., 2009), and through deacetylation of mitochondrial enzymes enhances metabolic flux of pathways often dysregulated in aging and age-related disorders (Kincaid and Bossy-Wetzel, 2013; McDonnell *et al*., 2015; Ansari *et al*., 2017). Here, we set out to investigate the importance of SIRT3 in CR-mediated longevity. We report that SIRT3 is dispensable for CR-dependent longevity, and unexpectedly, find that SIRT3 ablation further extends maximum lifespan under CR. Maximum lifespan is a parameter thought to correlate better with rates of aging as compared to average lifespan, since it is less likely to be influenced by strain specific diseases that can shorten life (Weindruch and Sohal, 1997). Though it is unclear if the extended maximum lifespan of *Sirt3^-/-^*CR mice is due to a retardation of the aging process, lifelong metabolic adaptations in *Sirt3^-/-^*CR mice may confer stress resistance late in life. SIRT3 deficiency attenuates succinate-dependent respiration but preserves palmitate-dependent respiration in heart and liver of aged CR-treated mice. *Sirt3^-/-^*CR mice were less able to maintain carbohydrate-derived energy production and displayed faster fuel switching to FAO during the postprandial period, leading to a longer fasting state relative to WTCR animals (Fig. 5). Despite longer maximum lifespan, altered fuel utilization in *Sirt3^-/-^*CR mice is associated with reduced exercise capacity and spontaneous activity.

**Fig 5.**
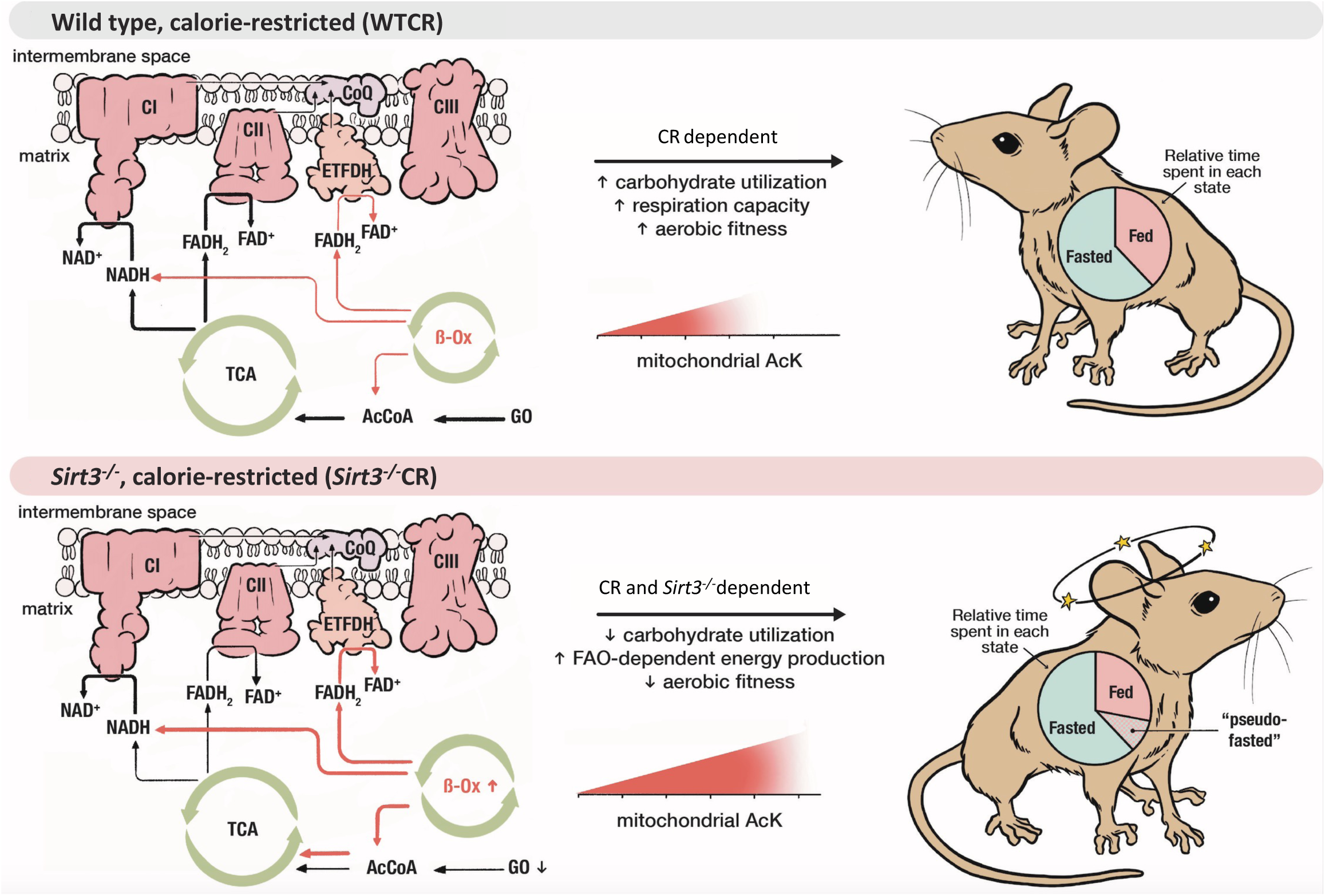
**Working model** Under fed state, energy is predominantly produced by glucose oxidation. However, due to *Sirt3^-/-^*-induced hyperacetylation, respiration capacity and TCA cycle efficiency are reduced, which limit the glucose oxidation-dependent energy production in *Sirt3^-/-^*CR mice and promote a fuel switching to fatty acid to compensate the energy demand that could not be met through glucose oxidation. This early shift to fatty acid oxidation produces a “pseudo-fasting” state, that is an increased fat oxidation period during the fed state. Longer fasting time has been found to promote lifespan extension in the mouse model. This prolonged fasting may explain the increased lifespan observed in *Sirt3^-/-^*CR mice. Despite increased lifespan, *Sirt3^-/-^*CR mice are less aerobically fit compared to WTCR mice due to inability of full glucose utilization, emphasizing the role of SIRT3 in preserving aerobic capacity.

Managing ROS production and detoxification has been proposed as a major SIRT3 function (Bause and Haigis, 2013; Zeng *et al*., 2014; Liu *et al*., 2017). In the current study, we found little evidence that Sirt3 deletion leads to ROS-dependent dysfunction. It is important to note that most prior aging and SIRT3 studies used C57BL/6J mice that lack functional NNT. The present study was conducted using the C57BL/6J/*Nnt^+/+^* genetic background. Likely, loss of both SIRT3 and NNT generated a more severe mitochondrial phenotype in which levels of NADPH and reduced glutathione (GSH) were significantly reduced (Qiu *et al*., 2010; Someya *et al*., 2010). We found that in the presence of functional NNT, NADPH and GSH levels were maintained in all treatment groups, strongly suggesting that the aging and metabolic phenotypes in the current study are independent of oxidative stress. Another important difference in our experimental design is the choice of a control diet that provides a fixed amount of 15% less calories as compared to ad libitum food intake. This control diet minimizes obesity-induced complications and allows a better comparison of the health benefits of CR relative to a non-pathological diet (Pugh, Klopp and Weindruch, 1999).

Loss of Sirt3 leads to a composite set of acetylation trends on mitochondrial proteins. While CR, age and loss of Sirt3 combined led to more sites of hyperacetylation, a detailed clustering analysis revealed sub-groups of sites in metabolic proteins that were dependent on both CR and Sirt3, but displayed decreased acetylation in *Sirt3^-/-^*CR mice. Fasting and CR increase protein acetylation in a tissue dependent manner (Schwer *et al*., 2009; Dittenhafer-Reed *et al*., 2015), likely by promoting FAO that increases mitochondrial acetyl-CoA production and consequently protein acetylation (Pougovkina *et al*., 2014; Mezhnina *et al*., 2020). Aging was also associated with elevated mitochondrial acetylation in various tissues (Joseph *et al*., 2012; Zeng *et al*., 2014). In current study, CR, aging and SIRT3 ablation increase overall liver mitochondrial acetylation individually and cooperatively, with proteins from oxidative metabolism pathways primarily affected, supporting the idea that SIRT3 counteracts a rise in protein acetylation in a subset of mitochondrial proteins under metabolic stress. KEGG pathway enrichment analysis of hyperacetylated proteins in old *Sirt3^-/-^* CR mice demonstrated that BCAA degradation and fatty acid degradation were highly enriched within this particular subset. Some of the BACC pathway acetylation sites with the largest increase in acetylation (Table S2) in the old *Sirt3^-/-^*CR mice compared to the baseline condition, the same age WTCD mice, included ACAD8 K220 (37% increase), ACAT1 K178 (16% increase), AUH K186 (10% increase), and ALDH7A1 K462 (50% increase). These sites were of particular interest because structural analyses indicate these sites are in close proximity to active sites or dimerization interfaces. For example, ACAD8 K220, ACAD8 K178, and AUH K186 are all adjacent to nearby NAD+, FAD+, and enoyl-CoA binding pockets, respectively (Kurimoto *et al*., 2001; Battaile *et al*., 2004, PDB2F2S). ALDH7A1 is a homotetramer, and K462 is found at the interface between the dimers of dimers. Others have suggested that NAD+ binding promotes the tetramerization of ALDH7A1 and its activation, therefore acetylation at this interface could be an important regulator of ALDH7A1 dehydrogenase activity (Luo *et al*., 2015; Korasick *et al*., 2018). Recent studies (Richardson *et al*., 2021; Yu *et al*., 2021) have shown that manipulation of BCAA level and its metabolism may lead to profound impact on metabolic parameters, including glucose control, adiposity, and longevity. Future studies are needed to determine the molecular and phenotypic consequence of SIRT3 loss in BCAA metabolism to resolve the influence of SIRT3 on aging mechanisms.

Unexpectedly, we identified a subgroup of acetylation sites that showed hypoacetylation in SIRT3 deficient animals on the CR diet. Though regulated by SIRT3 under CR, these sites are unlikely to be direct deacetylation substrates of SIRT3. The acetylation status of these sites could reflect the metabolic adaptation due to loss of SIRT3, leading to the unique phenotypes observed in *Sirt3^-/-^*CR mice. It is intriguing to speculate that acetylation of these sites is linked to acetyl-CoA availability, which could be positively regulated by SIRT3, for example, through PDH (Jing *et al*., 2013). Loss of SIRT3 leads to downregulation of PDH activity and reduced protein acetylation of these acetyl-CoA-sensitive sites, which include the site K304 on ACAT1 that was one of the most reactive lysine residues during non-enzymatic acetylation with acetyl-CoA (Baeza, Smallegan and Denu, 2016). Consistent with this idea, *Sirt3^-/-^*CR compared to WTCR mice show decreased levels of acetyl-CoA and citrate. Among this group of hypoacetylated sites in the *Sirt3^-/-^*CR condition, additional FAO pathway proteins include HADHA, ACAA2 and ECI1. Such protein acetylation sites may serve as an acetyl-CoA sensor and contribute to the FAO metabolic adaptation observed in *Sirt3^-/-^*CR mice.

Manipulation of SIRT3 expression results in clear changes in respiration phenotypes, with significant differences observed between the CR treated groups. SIRT3 ablation blunted CR-mediated respiration preservation and limited net OXPHOS respiration in *Sirt3^-/-^*CR animals, suggesting that both CR and SIRT3 are needed to maintain maximal bioenergetic capacity in aged mice. Notably, this CR and SIRT3- dependent respiratory phenotype is tissue dependent. Liver and heart displayed additive effects, whereas the brain showed only a genotype effect, and muscle showed neither genotype nor diet-induced effects. We also identified Complex II as the major contributor of reduced respiration in *Sirt3^-/-^*CR mice, consistent with studies (Cimen *et al*., 2010; Finley *et al*., 2011; Horton *et al*., 2016) showing that SIRT3 directly targets Complex II and restores activity through deacetylation. Several studies have also reported reduced Complex I activity in response to SIRT3 deletion (Ahn *et al*., 2008; Lantier *et al*., 2015; Williams *et al*., 2019). Although Complex I displayed altered function, compared to Complex II, Complex I activity and NADH-linked respiration were less affected in all tissues tested, suggesting Complex II dependent respiration is more prone to dysfunction under current experimental settings. Intriguingly, despite significant reduction in respiration (heart and liver) from *Sirt3^-/-^*CR animals using carbohydrate-derived substrates (pyruvate/malate/succinate), these mice maintained comparable palmitoylcarnitine/malate-dependent coupled respiration, indicating FAO is not limited. Recent studies (Lantier *et al*., 2015; Williams *et al*., 2019) report an enhanced palmitoylcarnitine/malate-dependent respiration phenotype in muscle of high-fat diet fed *Sirt3^-/-^* mice. While these are dramatically different dietary models (high fat diet vs. CR), the findings demonstrate that *Sirt3^-/-^*mice can preserve FAO capacity under both diets despite altered mitochondrial acetylation. Together, maintained/enhanced FAO respiration under CR or a high fat diet, and the composite trends in acetylation challenge the generalized idea that mitochondrial hyperacetylation limits FAO (Hirschey *et al*., 2010; Hallows *et al*., 2011; Tsuda *et al*., 2018), and instead highlights the importance of the dietary regime in the context of SIRT3-dependent mitochondrial regulation.

*Sirt3^-/-^*CR mice displayed faster fuel switching from glucose to FA compared to WTCR animals during the postprandial period. This was evident in RER and amount of FAO (Fig.4A-F) showing a trend of increased FAO during the postprandial period in *Sirt3^-/-^*CR relative to WTCR mice, in part due to limited Complex II function and partial redirected carbon flux to alanine and asparagine, indicating reduced glucose oxidation-dependent energy production. Unlike glucose oxidation, FAO requires an extra ETC component, the flavoprotein-ubiquinone oxidoreductase (ETF-QO, or ETFDH), to oxidize FADH_2_ harvested from beta-oxidation (Goetzman, 2011; Gnaiger *et al*., 2020). During acyl-CoA chain-shortening cycles, NADH and FADH_2_ are produced and further oxidized by Complex I and ETFDH, respectively. Switching to FAO may lead to more ETFDH- dependent electron transfer rather than transfer through Complex II. A recent study (Kľučková *et al*., 2020) reported that SDH-deficient murine chromaffin cells could maintain efficient FAO respiration that are higher than wildtype cells when palmitoyl-carnitine is respired. With only minorly affected Complex I observed in our study, FAO could be preserved in *Sirt3^-/-^*CR mice by increased flux through ETFDH, serving as a compensatory mechanism to drive FAO-dependent energy generation. Future studies are needed to determine whether compromised Complex II plays a role in fuel switching.

The early shift to FAO may lead to a “pseudo-fasting” state (Fig.5), which we define as increased fat oxidation period during the fed state. This pseudo-fasting state contributes to a longer overall FAO time and ultimately a longer fasting time in *Sirt3^-/-^*CR compared to WTCR mice. Increased fasting period has been a feature of meal-fed CR models, in which CR animals quickly consume their food allotment and fast until the next meal (Acosta-Rodríguez *et al*., 2017). Intriguingly, Pak *et al*. (2021) found that even without CR, increasing fasting time alone can promote CR-like metabolic phenotypes, highlighting the necessity of fasting towards CR benefits. Both alternate day feeding and daily fasting increase longevity in mice and the beneficial effects of these interventions have been shown to be independent of caloric intake (Mattson *et al*., 2014; Mitchell *et al*., 2019). Fasting is also associated with improved intestinal stem cell function in aging (Mihaylova *et al*., 2018). Interestingly, the long-lived Ames dwarf and GHR-knockout mice display increased reliance on fatty acid oxidation as a fuel source (Bartke and Westbrook, 2012). Thus, existing studies have linked increased fasting or increased fatty acid oxidation with health benefits in rodents. We speculate that this additional pseudo-fasting period may play an important role in extending the maximum lifespan of *Sirt3^-/-^*CR mice. Notably, the feeding protocol used in the current study involves an every-other-day feeding schedule for both CD and CR animals. This design increases fasting time between feedings and provides a more rigorous examination on the effects of a CR regime, as compared to studies that use ad libitum feeding for controls. Despite increased lifespan, *Sirt3^-/-^*CR mice are less aerobically fit compared to WTCR mice due to inability of full glucose utilization (Seiler *et al*., 2015), emphasizing the role of SIRT3 in preserving aerobic capacity.

Taken together, our findings reveal that SIRT3 is required for optimal aerobic fitness during aging but not for CR-mediated longevity. These results highlight the uncoupling of lifespan and healthspan parameters such as aerobic fitness and spontaneous physical activity, and address the need to comprehensively assess the factors that contribute to CR-dependent phenotypes. Possibly, some CR-induced factors contribute to lifespan extension, while having no impact on healthspan, and vice versa.

## AUTHOR CONTRIBUTIONS

Conceptualization, J.M.D., and T.A.P.; Methodology, R.S.D., Y.Q., F.B.M., C.A.G., D.W.L., T.A.P., and J.M.D.; Investigation, R.S.D., Y.Q., P.R.G., V.X.F., A.J.L., C.L.G., F.B.M., and C.A.G.; Resources, J.M.D., T.A.P., D.W.L., F.B.M., and C.A.G.; Animal breeding and care, V.X.F., and J.M.V.; Writing-Original Draft, Y.Q., J.M.D., and T.A.P.; Writing-Review & Editing: Y.Q, R.S.D, P.R.G, F.B.M., C.A.G., D.W.L., J.M.D., and T.A.P.; Funding Acquisition, J.M.D., T.A.P., D.W.L., F.B.M., and C.A.G.,

## ACKNOWLEDGMENTS

This work was supported by NIA grant AG038679 and GM65386 to T.A.P and J.M.D. The Lamming laboratory is supported in part by the NIH/National Institute on Aging (AG056771, AG062328, and AG061635 to D.W.L.) and startup funds from the University of Wisconsin-Madison School of Medicine and Public Health and Department of Medicine to D.W.L. C.L.G. is supported by a Glenn/AFAR Postdoctoral Fellowship from the Glenn Foundation for Medical Research. Support for this research was provided by the University of Wisconsin - Madison Office of the Vice Chancellor for Research and Graduate Education with funding from the Wisconsin Alumni Research Foundation. The Lamming lab is supported in part by the U.S. Department of Veterans Affairs (I01-BX004031), and this work was supported using facilities and resources from the William S. Middleton Memorial Veterans Hospital. The content is solely the responsibility of the authors and does not necessarily represent the official views of the NIH. This work does not represent the views of the Department of Veterans Affairs or the United States Government. F.B.M. and C.A.G. were supported by FAPESP (# 2015/00272-6, # 2015/01362-9) for their working period at University of Wisconsin-Madison. We would like to thank Randall Massey at the Medical School Electron Microscope Facility for his help in transmission electron microscopy. We thank Dr. Melissa Skala, Dr. Alex Walsh, and Kelsey Tweed for the insightful discussions and comments. We also thank Eric Armstrong for his help for metabolomics.

## DATA AND CODE AVAILABILITY

The raw data, processed data, spectral library, and the analysis logs describing the settings for the Spectronaut analyses have been deposited to the ProteomeXchange Consortium via the MassIVE partner repository with the dataset identifier MSV000087085 and PXD024961 [doi:10.25345/C59Z2Q]. The data was processed and cleaned using an in-house R script, which can be accessed through the GitHub link: [DOI:10.5281/zenodo.3360892]. Other data are available upon request.

## EXPERIMENTAL MODEL AND SUBJECT DETAILS

All animal studies were conducted at the AAALAC-approved Animal Facilities at the William S. Middleton Memorial Veterans Administration Medical Center and the University of Wisconsin-Madison under animal research protocols approved by these institutions Institutional Animal Care and Use Committee (IACUC).

### Animals

Male and female *Sirt3^+/−^* mice (Stock #011664-UNC) were purchased from the Mutant Mouse Resource Centers (MMRRC) at the University of North Carolina-Chapel Hill (Chapel Hill, NC). These mice were created by a retroviral promoter trap that functionally inactivates one allele of the Sirt3 gene by a 5.1 kb retroviral insertion in the intron preceding coding exon 1 (MGI:3529767). *Sirt3^+/+^* C57BL/6J mice were purchased from The Jackson Laboratory (Stock number 000664). *Sirt3^+/−^* mice were then backcrossed onto the C57BL/6J background (Someya *et al*., 2010). To obtain the functional NNT, *Sirt3^-/-^* C57BL/6J mice were interbred with *Sirt3^+/+^* C57BL/6N (*Nnt^+/+^*) mice (The Jackson Laboratory, stock number 005304). Interbreeding produced C57BL/6J/*Nnt^+/+^ Sirt3^+/-^* mice were further interbred to produce the two strains of mice used in experiments: C57BL/6J/*Nnt^+/+^ Sirt3^-/-^* mice and their wild-type counterparts C57BL/6J/*Nnt^+/+^ Sirt3^+/+^*mice.

### Genotyping

Genotyping for *Nnt* and *Sirt3* were accomplished using PCR reactions. DNA was prepared from tail snips obtained as mice were weaned at 3 weeks of age. The *Sirt3* genotyping reaction consisted of final concentrations of 1X reaction buffer and 0.05 U/ul Taq polymerase (Denville, Metuchen, NJ), 0.20 µM each primer (IDT), 0.20 mM dNTP mix and 1.75 mM MgCl_2_ (Thermo-Fisher/Invitrogen). *Sirt3^+/+^* mice were indicated by a 404 bp product across the insertion points of the (absent) viral insert. *Sirt3^-/-^*mice were indicated by two amplification products: 207 bp across the 5’ insertion site of the (present) viral insert and 291 bp across the 3’ insertion site. Heterozygous mice were indicated by the presence of all three bands. The *Nnt* genotyping reaction consisted of final concentrations of 1X reaction buffer and 0.10 U/ul Taq polymerase (Denville, Metuchen, NJ), 0.40 µM each primer (IDT), and 0.20 mM dNTP mix (Thermo-Fisher/Invitrogen). *Nnt^+/+^* mice were indicated by a 304 bp product across the deletion point of the mutation, which confirmed there was no deletion. *Nnt^-/-^* mice were indicated by two amplification products: 227 bp across the 5’ deletion site and 203 bp across the 3’ deletion site. Heterozygous mice were indicated by the presence of all three bands.

### Husbandry and Diets

Starting at two months of age, male *Sirt3^-/-^*and *Sirt3^+/+^* mice were individually housed, randomly assigned to a control diet (CD) or calorie restricted diet (CR). Both CD and CR mice maintained their assigned diet till death. The control group was fed 96 kcal/week of diet AIN-M (Bio-Serve, Farmington, NJ), which is ∼15% less than the average ad lib intake (Pugh, Klopp and Weindruch, 1999). The CR group was fed 72 kcal/week. The restricted diet food pellets are a modified form of AIN-M (Bio-Serve) and are nearly isocaloric gram-for-gram with the control diet food pellets, but enriched in protein, vitamins and minerals to provide the same amounts of these components as control diet mice receive. CD and CR mice were fed on Monday/ Wednesday/Friday at approximately 7:00 a.m. (CD mice: 7 grams on Monday/Wednesday, and 10 grams on Friday; CR mice: 5 grams on Monday/Wednesday, and 8 grams on Friday).

Any mice that appear ill, cachectic, stressed, less than 22 grams indicated by cage card were euthanized. As these mice age, further criteria for euthanasia included: body weight loss of 20% in one month, and persistent severe ulcerative dermatitis. Mice that have a body condition score less than 1.5 on the Ullman-Cullere and Foltz Body Condition Scale (Ullman-Culleré and Foltz, 1999) were euthanized.

### Survival Study

All animals were inspected each day. Mortality during the survival study was analyzed using the log-rank test to compare the differences in Kaplan-Meier survival curves. Median lifespan and maximal lifespan were defined as the time of 50% mice survival and the average lifespan of top 10% longest lived mice. Maximum lifespan comparison was performed using two-way ANOVA followed by multiple t-test, corrected by Tukey’s test. Maximum lifespan statistics was also confirmed with Boschloo’s Test. Graph Pad Prism 9.1.1 was used for visualization and OASIS-2 (Han *et al*., 2016) was used for Log-rank test and Boschloo’s Test.

### Body Weight and Composition

Two-month-old mice, immediately after entering CR, were weighed weekly. Exceptions were made when weight decreased by one gram (or ∼5% body weight) in one week, in which case the mice were weighed on the same days as they were fed until they gained weight. If a mouse lost 4 grams (or ∼20% body weight) in one week, it was euthanized. Adult CR mice (>6 months) were weighed weekly as long as their weight was at or above 25 grams. If their weight was between 22-25 grams, they were then weighed on the same days as feeding. If mice weighed less than 22 grams, or their body condition score decreased to 2.0 or less, they were weighed daily. The control group of mice were weighed once per month. Control mice were then weighed once every two weeks starting at age 24 months and weighed weekly staring at age 29 months (or 50% mortality rate was reached). Mouse body composition was determined using the EchoMRI Body Composition Analyzer (EchoMRI, Houston, TX, USA).

### Transmission Electron Microscopy

Tissue for transmission electron microscopy (TEM) was sampled from muscle fibers collected from the gastrocnemius and heart left ventricular wall. Tissue samples were fixed and mounted at the Medical School Electron Microscope Facility (University of Wisconsin-Madison). Briefly, samples were fixed in 2.5% glutaraldehyde in 0.1 M phosphate buffer, post-fixed in 1.0% osmium tetroxide, and dehydrated in a graded series of ethanol concentrations (30-100%), prior to embedding in epoxy resin. Ultrathin (0.5 µm) sections were then mounted for post fixing in uranyl acetate and lead citrate. Sections were viewed using a Philips CM120 transmission electron microscope and images were captured with a BioSprint 12 series digital camera. Four mice per treatment group at 25 months of age were analyzed. Measurements were taken at a final magnification of 5600x and 19500x for mitochondrial volume density and morphology, respectively. Volume density was calculated using the summed point count method (Schmiedl *et al*., 1990). This method uses the number of intersections of a grid landing on mitochondria relative to those intersections landing on the reference space as an index of volume density. TEM samples were analyzed blindly by preparing samples with a new set of numbers and cross referenced after analysis to remove any experimenter bias during the measurement process.

### Tissue Homogenate Preparation

Pulverized tissue was suspended in hypotonic buffer (25 mM K_2_HPO4, 5 mM MgCl_2_, pH 7.2) with protease inhibitor cocktail (100X, Thermo Scientific). The mixture was then sonicated for three 10-second cycles (10 seconds pauses between each cycle, 30% amplitude) and centrifuged at 1,000xg for 10 mins at 4°C. Supernatant was transferred to a new tube and centrifuged again at 1,000xg for 10 mins at 4°C. Total protein concentration of the supernatant was determined using Pierce BCA protein assay kit (Thermo Scientific). Supernatant was aliquoted and stored at -80°C.

### Citrate Synthase Activity

Citrate synthase activity assay was described by Hepple *et al*., (2005). Briefly, citrate synthase activity was determined by measuring the production rate of TNB (thionitrobenzoic acid) at an absorbance of 412nm. To initiate the reaction, reaction buffer (final concentration: 0.3mM acetyl-CoA, 100mM Tris buffer pH 8.0, 0.1mM DTNB, and 0.5mM oxaloacetate) was added into properly diluted tissue homogenate. Absorbance at 412nm was measured using a plate reader (Synergy™ H4 Hybrid Multi-Mode Microplate Reader) at 37°C. Citrate synthase activity was normalized to the amount of total protein in each sample.

### Mitochondrial DNA Content

Methods to determine the copy number of mitochondrial DNA per nucleus is described by Patil *et al*., (2015). Intronic sequences close to those used by Patil *et al*., (2015) for LPL (NC_000074.6) and Nd1 (NC_005089.1). A multiplex PCR was designed using a FAM-labeled probe for LPL and a HEX-labeled probe for Nd1. Both probes incorporated the internal ZEN™ dark quencher in addition to the 3’ quencher Iowa Black^®^ FQ.

A gBlocks® Gene Fragment (IDT) served as a standard curve template for both LPL and Nd1 to determine copy number. The oligonucleotide template includes 246 bp of the LPL sequence and 213 bp of the Nd1 sequence. Standards included with each PCR run were 0, 10, 100, 1,000, 10,000, 33,000, 100,000, and 1 million copies of the template. RNA and DNA were prepared by an adaptation of the method of Hofer *et al*., (2006). Approximately 50 mg of tissue was homogenized using a Tissuelyzer II (Qiagen) at 25 cycles/second for two runs of 30 seconds each in GTC buffer (3M guanidine thiocyanate, 0.2%lauroylsarcosinate, 20 mM Tris buffer, pH 7.5) made to 20 mM deferoxamine mesylate immediately before use. An equal volume of phenol-chloroform-isoamyl alcohol (25:24:1) pH 6.7 was added, vortexed immediately and repeatedly vortexed during a 10-minute incubation at room temperature. Samples were centrifuged at 14,000xg at 4°C for 10 min. Then the upper aqueous layer was removed, mixed with an equal volume of isopropanol, and precipitated at -20°C for one hour. Samples were then centrifuged at 14,000xg at 4°C for 10 min and washed twice with 70% ethanol. The pellet containing RNA and DNA was dried and re-dissolved in 50 µl of chilled nuclease-free water. DNA was separated from RNA using the Macherey-Nagel Nucleospin Tri-prep spin column kit according to the manufacturer’s instructions (Macherey-Nagel, Düren, Germany). RNA and DNA were then re-precipitated in 70% ethanol, 0.83 M sodium acetate and washed twice with 70% ethanol. Samples were re-dissolved in 50 µl of TE buffer. After initial reading of DNA concentrations in a Nanodrop 2000 (Thermo-Fisher) samples were diluted to approximately 7.5 ng/µl. Concentrations were measured again using a Quant-IT Fluorescent assay (Thermo-Fisher). These final concentration values were used to normalize the PCR results.

PCR reactions consisted of PrimeTime Gene Expression Master Mix (IDT), LPL primers and FAM-labelled probe, Nd1 primers and HEX-labelled probe and 2 µl of sample containing approximately 15 ng of sample DNA in a final reaction volume of 10 µl. Real-time PCR was performed in an Eppendorf Realplex Thermocycler with an initial denaturation step of 95°C for 3 min, then 40 cycles of 95°C for 5 seconds and 60°C for 30 seconds. Cycle thresholds of the standards were used to generate a standard curve to calculate the copies of mtDNA and genomic DNA in each sample.

### Automated Capillary Electrophoresis Based Immunoblot

All proteins analyzed by automated capillary electrophoresis-based immunodetection system (WES, ProteinSimple) were detected using the 12-230kDa WES separation module (SM-W004, ProteinSimple) and a detection module (depending on primary antibody source, an anti-rabbit (DM-001) or anti-mouse (DM-002) detection module was used). Briefly, tissue lysates in RIPA buffer were diluted in 0.1x sample buffer (optimized per tissue in preliminary experiments) and 8µl was combined with 2µl 5x fluorescent master mix. Samples were denatured by heating at 95°C for 5 minutes (50°C, 5 minutes for detection of ETC proteins). For immunodetection, blocking buffer, primary antibody (in per tissue optimized dilution), secondary antibody, chemiluminescent substrate, samples (including a biotinylated size marker) and wash buffer were loaded in in the above order in the designated wells on the supplied microplate. The plate was centrifuged for 5 mins at 1,000xg to remove air bubbles and loaded into WES. Default separation parameters were used for single protein detection. For ETC Complex IV and multiplexing of ETC Complex III and V specific antibodies, sample loading time and separation time were increased to 21mins and 31mins, respectively. Following automated protein separation and detection, data were analyzed using Compass for SW software (version 3.1.7, ProteinSimple) and the area under the specific antibody peak was measured. To compare protein expression among samples of a tissue type, expression was normalized to total protein content. Total protein was analyzed in all the samples in a separate set of assays using a biotin-based protein labeling and detection system (DM-RP01) on the WES. Total protein was also analyzed using Compass for SW software as total area under the peaks. Two major peaks that were determined to result from direct binding of the biotin detection reagent to protein were excluded from analysis. Control carryover samples between plates were used to correct for inter-plate signal intensity variability (generally not exceeding 10%). The area under the specific antibody peak in each sample was divided by the area under the total protein peaks of that sample and averaged among the same tissue samples from the mice of the same genotype on the same diet (n=6). Primary antibodies were used to detect Citrate Synthase (#14309, CST), SIRT3 (#5490, CST), NDUFA9 (ab14713, abcam), SDHB (ab14714, abcam), UQCRC2 (ab14745, abcam), MTCOI (ab14705, abcam) and ATP5A (ab14748, abcam).

### Mitochondria Isolation

Freshly harvested tissue was quickly rinsed in 1xPBS and placed in an ice-cold petri-dish with isolation media (250mM sucrose, 25 mM KH_2_PO_4_, 50 mM KCl, 10 mM HEPES, 0.5 mM EGTA, 1% BSA, pH 7.4). Tissue was finely minced and transferred to an ice-cold glass homogenizer. Tissue was homogenized with a tight pestle on ice. The number of strokes was optimized for each tissue. Homogenate was then transferred into a conical tube and centrifuged at 900xg for 10 mins at 4°C. Supernatant was filtered through a layer of glass wool and centrifuged at 9,000xg for 10 mins at 4°C. Supernatant was discard, and the mitochondrial pallet was resuspended in isolation media before being centrifuged again at 9,500xg for 10 mins at 4°C. The supernatant was again discarded, and the mitochondrial pellet was suspended in storage buffer (25 mM KH_2_PO_4_, 50 mM KCl, 10 mM HEPES, 0.5 mM EGTA, pH 7.4) and stored at -80°C.

### Western Blot

Pulverized tissues or isolated mitochondria were lysed in RIPA buffer (50mM Tris-HCl, 150mM NaCl, 0.1% Triton-100, 0.5% sodium deoxycholate, 0.1% SDS, 1mM EDTA, protease inhibitor cocktail (100X, Thermo Scientific), 10mM nicotinamide and 10mM sodium butyrate). Tissue lysates were sonicated for three 5-second cycles (10 seconds pauses between each cycle, 20% amplitude) and mitochondrial lysates were agitated for 1 hours at 4°C. Lysates were centrifuged at 12,000rpm for 20 mins at 4°C, and the supernatants were placed into new tubes. Total protein concentration of each sample was determined by Pierce BCA protein assay kit (Thermo Scientific). Each sample, containing ∼30µg total protein, was mixed with loading buffer (4X, LI-COR) and heated for 5 mins at 95°C. These samples were separated on 10% in-house casted SDS-Page gels and transferred onto nitrocellulose membrane (GE). Total protein stain (LI-COR) was used for total protein normalization. After washing for 10 mins in TBST (20mM Tris, pH7.5, 150mM NaCl, 0.1% Tween-20), membranes were blotted in 5% BSA in TBST at room temperature for 1 hour and incubated overnight with primary antibody (in 2% BSA TBST) solution at 4°C. The next day, membranes were washed with TBST and incubated with secondary antibody (LI-COR, 1:10,000) for 90 mins at room temperature. Membranes were washed for 5 minutes in TBST 3 times and followed by two 5-min TBS wash. Membranes were imaged using the Odyssey scanner (LI-COR Odyssey). Images were further analyzed by Image Studio Lite version 5.2.5 (LI-COR). Western blot primary antibodies used include: SIRT3 (#5490, CST, 1:1000), Acetylated-Lysine (#9681, CST, 1:1000), VDAC (75-204, NeuroMab, 1:1250).

### LC-MS Based Quantification of Mitochondrial Acetylation Stoichiometry

Quantification of acetylation stoichiometry follows methods described previously in Baeza *et al*., 2020, with the following modifications. Notably, the current study quantified acetylation stoichiometry of individual lysine residues using an antibody free approach, which does not involve acetyl-peptide immunoprecipitation step during sample preparation. Analysis was conducted using 200 μg of protein from isolated mitochondrial subcellular fractions, which were denatured in urea buffer (6 M urea (deionized), 100 mM ammonium bicarbonate pH = 8.0, 5 mM DTT). Samples were incubated for 20 minutes at 60 °C, then cysteines were alkylated with 50 mM iodoacetamide and incubated for 20 minutes. Chemical acetylation using two rounds of ∼20 µmol heavy isotopic D6-acetic anhydride (Cambridge Isotope Laboratories). Samples were diluted using 100 mM ammonium bicarbonate pH = 8.0 to 2 M urea and digested with 1:100 trypsin at 37 °C for 4 hours. Samples were then diluted to 1 M urea prior to a second digestion by gluC (1:100). Chemically acetylated peptides were fractionated into 6 fractions using a Shimadzu LC-20AT HPLC system with a Phenomenex Gemini^®^ NX-C18 column (5µm, 110Å, 150 x 2.0mm). The samples were analyzed using data-independent acquisition (DIA) analysis by a Thermo Q-Exactive Orbitrap coupled to a Dionex Ultimate 3000 RSLC nano UPLC with a Waters Atlantic reverse phase column (100 μm x 150 mm).

To deconvolute and analyze the DIA spectra, a spectral library containing all light and heavy acetyl-lysine feature pairs was generated. Spectral library samples were processed identically to the experimental samples, except that they were treated with C12-acetic anhydride (Sigma) and analyzed using data dependent acquisition (DDA) mass spectrometry analysis. A spectral library was generated using the openly available MaxQuant (v1.6.0) software package. Carbamidomethylation (C) was set as a fixed modification, and Oxidation (M) and Acetyl (K) were set as variable modifications. Trypsin and gluC were set as the digestion enzymes, with the max number of missed cleavages set to five. DDA runs from both the mitochondrial and cytosolic fractions were run to make one combined library. Heavy acetyl fragment ion pairs were generated in silico, such that the spectral library would contain both the light (endogenous) acetylation peaks and the heavy (chemical) acetylation peaks. The experimental samples were processed using Spectronaut (v10) using the generated spectral library. The subcellular fraction experimental samples were processed separately. The data was processed and cleaned using an in-house R script, which can be accessed through the GitHub link: [DOI:10.5281/zenodo.3360892], such that stoichiometry was calculated from the ratio of endogenous (light) fragment ion peak area over the total (endogenous and chemical) fragment ion peak area. One-way ANOVA analyses were used to determine statistically significant differences in acetylation stoichiometry driven by age, diet, or genotype between two groups. The raw data, processed data, spectral library, and the analysis logs describing the settings for the Spectronaut analyses have been deposited to the ProteomeXchange Consortium via the MassIVE partner repository with the dataset identifier MSV000087085 and PXD024961 [doi:10.25345/C59Z2Q].

### Critical Velocity

The critical velocity (CV) was determined according to the protocol previously described (Scariot *et al*., 2019), based on the mathematical relationship between exercise intensity and time to exhaustion (tlim). Briefly, animals were subjected to four exhaustive running efforts applied on different days, with the time to exhaustion recorded for each exercise intensity. Before the actual experiment, mice were adapted to a treadmill ergometer (Exerc 3/6 Treadmill, Columbus Instruments, Ohio, USA) and to exercise for three days (5 min per day). To determine CV, constant exercise intensities (treadmill velocity ranging from 7.5 to 20.0 m/min) were individually selected so that the time to exhaustion was no more than 15 min and no less than 1 min. The time to exhaustion, recorded in seconds, was determined when the mouse was unable to run despite encouragement (gentle soft brush tapping, without electrical stimulus). Distance traveled for each mouse during individual tests was calculated by multiplying the velocity of the treadmill and time to exhaustion. Distance traveled and time to exhaustion, total four data points, were then plotted and fitted into a linear regression model. The goodness of the linear regression was evaluated by R^2^. The slope of the fitted regression corresponded to the critical velocity of the mouse.

### Mitochondrial Respiration in Permeabilized Tissues

Brain (cerebral cortex), gastrocnemius fiber, heart and liver were freshly harvested from mice sacrificed by acute conscious cervical dislocation on the day of respiration experiments and temporally stored in iced cold BIOPS buffer (2.77 mM CaK_2_EGTA, 7.23 mM K_2_EGTA, 50 mM MES hydrate, 0.5 mM dithiothreitol, 20 mM imidazole, 20 mM taurine, 15 mM sodium phosphocreatine, 6.56 mM MgCl2, 5.77 mM ATP). Tissues were then permeabilized in BIOPS containing 30µg/ml saponin for 30 mins at 4°C with gentle shaking (detailed tissue permeabilization steps were described in Doerrier *et al*., (2018) and Kuznetsov *et al*., (2008). After being permeabilized, tissues were placed in cold Mir05 buffer (110 mM sucrose, 60 mM potassium lactobionate, 0.5 mM EGTA, 3 mM MgCl_2_, 20 mM taurine, 10 mM KH_2_PO_4_, 20 mM HEPES, 1g/L BSA, pH 7.1) and stirred generally for 10 mins at 4°C. Wet tissue weight was then collected, and tissue was place into the chamber. Mitochondrial respiration was determined by measuring oxygen consumption rate (OCR) using OROBOROS O2k (Oroboros Instruments) in Mir05 buffer at 37°C. When oxygen consumption rate reached steady state, substrates were added in the following order with final concentration in 2ml chamber. 5mM pyruvate/ 2mM malate /10mM glutamate were added to establish leak respiration, followed by 1mM ADP to stimulate a coupled respiration rate. 10µM Cytochrome C was added to assess any outer membrane damage to the mitochondria. Then 10mM succinate was used to stimulate complex II-fueled respiration, followed by FCCP to measure uncoupled respiration. 0.5µM rotenone was used to isolate complex II respiration from complex I, and finally 2.5µM antimycin was used to estimate residual respiration. To examine fatty acid oxidation-dependent respiration, substrates were added in the following order with final concentration in 2ml chamber, 40 µM palmitoyl-carnitine with 0.1mM malate/ 1mM ADP/10µM Cytochrome C/ FCCP/2.5µM antimycin. Reported OCR for non-fatty acid substrates were normalized to CS activity, and OCR for non-fatty acid substrates were normalized to total protein.

### Mitochondrial Complex Activity

Complex I and Complex II activity were determined by measuring the reduction of DCPIP with an absorbance of 600nm. Complex I activity was measured in triplicate by adding 250µl reaction mix A (hypotonic medium (25mM K_2_HPO_4_, 5mM MgCl_2_, pH7.2), 2.5mg/ml BSA, 100µM DCPIP, 65µM ubiquinone-2, 2µg/ml antimycin, 0.2mM NADH) to 5µl of properly diluted tissue homogenate (dilution was optimized for each tissue). Background was measured by adding 250µl reaction mix A with 5µM rotenone, to tissue homogenate. The reaction was monitored at an absorbance of 600nm for 20 mins. Complex II activity was measured in triplicate by adding 250µl reaction mix B (hypotonic medium (25mM K_2_HPO_4_, 5mM MgCl_2_, pH7.2), 2.5mg/ml BSA, 100µM DCPIP, 65µM ubiquinone-2, 2µg/ml antimycin, 2µg/ml rotenone, 20mM succinate) to 5µl of properly diluted tissue homogenate. Background was measured by adding 250 µl reaction mixture B, without succinate, to tissue homogenate. The reaction was monitored at an absorbance of 600nm for 20 mins. Reported Complex I and II activity were corrected to Complex I and Complex II expression, respectively, which were measured by WES as described in earlier sections.

### Metabolite Extraction and LC-MS-based Metabolite Profiling

The metabolite extraction method was described in Haws *et al*. (2020) with the following modifications. Tissues were pulverized prior to metabolite extraction. Pulverized tissue (∼30mg) was incubated with 1ml of ice cold 80:20 methanol: water solution for 5 mins on dry ice after 15-second vortexing. Tissue homogenate was centrifuged at maximum speed for 5 minutes at 4°C. Supernatant was collected and transferred into a new tube. The remaining pellet was incubated with 400µl of ice cold 40:40:20 methanol:acetonitrile:water solution for 5 mins. Tissue homogenate was centrifuged again at maximum speed for 5 minutes at 4°C. Supernatant was pooled with the first metabolite extraction. The 40:40:20 methanol:acetonitrile:water solution extraction was repeated once more and combined with the previous extractions. Three extractions were pooled and completely dried using SpeedVac (Thermo Fisher Savant ISS110) with nitrogen flow under room temperature. The dried metabolite samples were resuspended in water (150µl water per 5mg tissue) and centrifuged at maximum speed for 5 minutes at 4°C. This supernatant was used for LC-MS metabolite profiling.

The metabolite detection method was adopted from Latorre-Muro *et al*. (2018). Briefly, metabolites were separated by Thermo Fisher Vanquish UHPLC with Waters Acquity UPLC BEH C18 column (1.7 μm, 2.1 × 100 mm; Waters Corp.) and analyzed by Thermo Fisher Q Exactive orbitrap mass spectrometer in negative ionization mode. LC separation was performed over a 25-minute method with a 14.5-minute linear gradient of mobile phase (buffer A, 97% water with 3% methanol, 10 mM tributylamine, and acetic acid-adjusted pH of 8.3) and organic phase (buffer B, 100% methanol) (0min, 5%B; 2.5min, 5%B; 17min, 95%B; 19.5min, 5%B; 20 min, 5%B; 25 min, 5%B, flow rate 0.2ml/min). 12µl of each sample was injected into the system for analysis. ESI settings were 30/10/1 for sheath/aux/sweep gas flow rates, 2.50 |kV| for spray voltage, 50 for S-lens RF level, 350°C for capillary temperature, and 300°C for aux gas heater temperature. MS1 scans were operated at resolution = 70,000, scan range = 85-1250 m/z, automatic gain control target = 1e6, and 100ms maximum IT. Metabolites were identified and quantified using El-MAVEN (v0.12.1-beta, Agrawal *et al*., 2019) with metabolite retention times empirically determined in-house. Metabolite levels were compared using the peak AreaTop.

### Spontaneous Physical Activity

The gravimetric method used to measure the spontaneous physical activity (SPA) was adopted from Biesiadecki *et al*. (1999) and Scariot *et al*. (2016) with modifications. In this study, the individual cage was put on the metallic platform equipped with one load cell (model PLA 10kgf, Lider Balanças, BR), capable of identifying the force generated by rodent activities. The instruments used to amplify and condition the signals were MKTC-05 (MK Controle e Instrumentação, BR) and NI-USB 6008 (National Instruments, USA), respectively. The signals were captured using digital acquisition software (LabVIEW SignalExpress National Instruments, USA) at 30Hz frequency. The gravimetric system was calibrated before the experiments by positioning known weights on the central point of the force platform. The signals obtained in volts were converted to units of gram using the linear regression equations (all calibrations presented R^2^ = 1.00) acquired from calibrations plots. Data were processed by using Matlab (R2008a MatLab, MathWork) as described in Scariot *et al*. (2019). In the current study, SPA, in arbitrary units, was registered during a 19-day period (including both light and dark phases, in all groups of 25-month-old mice).

### Quantitative PCR Analysis

RNA was first isolated from ∼30mg pulverized tissue using TRIzol Reagent (Invitrogen). Genomic DNA was removed using DNase I (Thermo Scientific) and cDNA was synthesized using RevertAid First Strand cDNA Synthesis Kit (Thermo Scientific) following manufacturer’s instructions. Then, the real-time PCR reactions were performed in triplicates using PerfeCTa SYBR Green SuperMix (Quantabio) and a Bio-Rad CFX96 C1000 Real-Time system. The expression levels of target genes were normalized against the housekeeping gene β2-microgulin.

### Whole Body Metabolic Assessment

To measure metabolic parameters [O2, CO2, food consumption, respiratory exchange ratio (RER), energy expenditure], mice were acclimated to housing in a Oxymax/CLAMS metabolic chamber system (Columbus Instruments) for ∼2 h and data from a continuous 48 h period was then recorded and analyzed. When cages were opened for addition/removal of food, data points were removed. RER, EE values were reported by the metabolic chamber system. FAO per hour were calculated by the equation: FAO (kcal/hr) = EE x (1-RER/0.3) (Bruss *et al*., 2010). When calculated FAO <0, 0 was assigned to FAO for that hour. 24 hours FAO (kcal/24hr) was calculated from the 24-hr area under the curve of hourly FAO. Reported 24 hours FAO per mouse was corrected to its body weight.

## QUANTIFICATION AND STATISTICAL ANALYSIS

Lifespan was assessed using the log-rank test by comparing the differences in Kaplan-Meier survival curves (Mitchell *et al*., 2019). Maximum lifespan was assessed by the average lifespan of the top 10% longest lived mice and Boschloo’s Test. Both Log-rank test and Boschloo’s test were conducted using Online Application for Survival Analysis 2 (OASIS 2) with default settings (Han *et al*., 2016). Results are plotted as mean ± SEM unless otherwise specified, with *p*≤0.05 considered statistically significant. Outliers were excluded using ROUT method with Q = 5%. Data were analyzed by one- or two-way ANOVA followed by t-test. *p* values reported for each comparison were corrected by Tukey HSD post-hoc test when it is applicable. Significant (p≤0.05) diet effect, genotype effect and/or diet and genotype interaction for each experiment are indicated in figures. Metabolite levels and gene expression were analyzed by unpaired t-test with *p* value uncorrected for multiple comparisons. Analyses and visualization were performed using Graph Pad Prism (9.0.0, GraphPad Software), Excel (Version 16.52, Microsoft), RStudio (Version 1.4.1106), Matlab (R2008a MatLab, MathWork).

---

**Fig S1.**
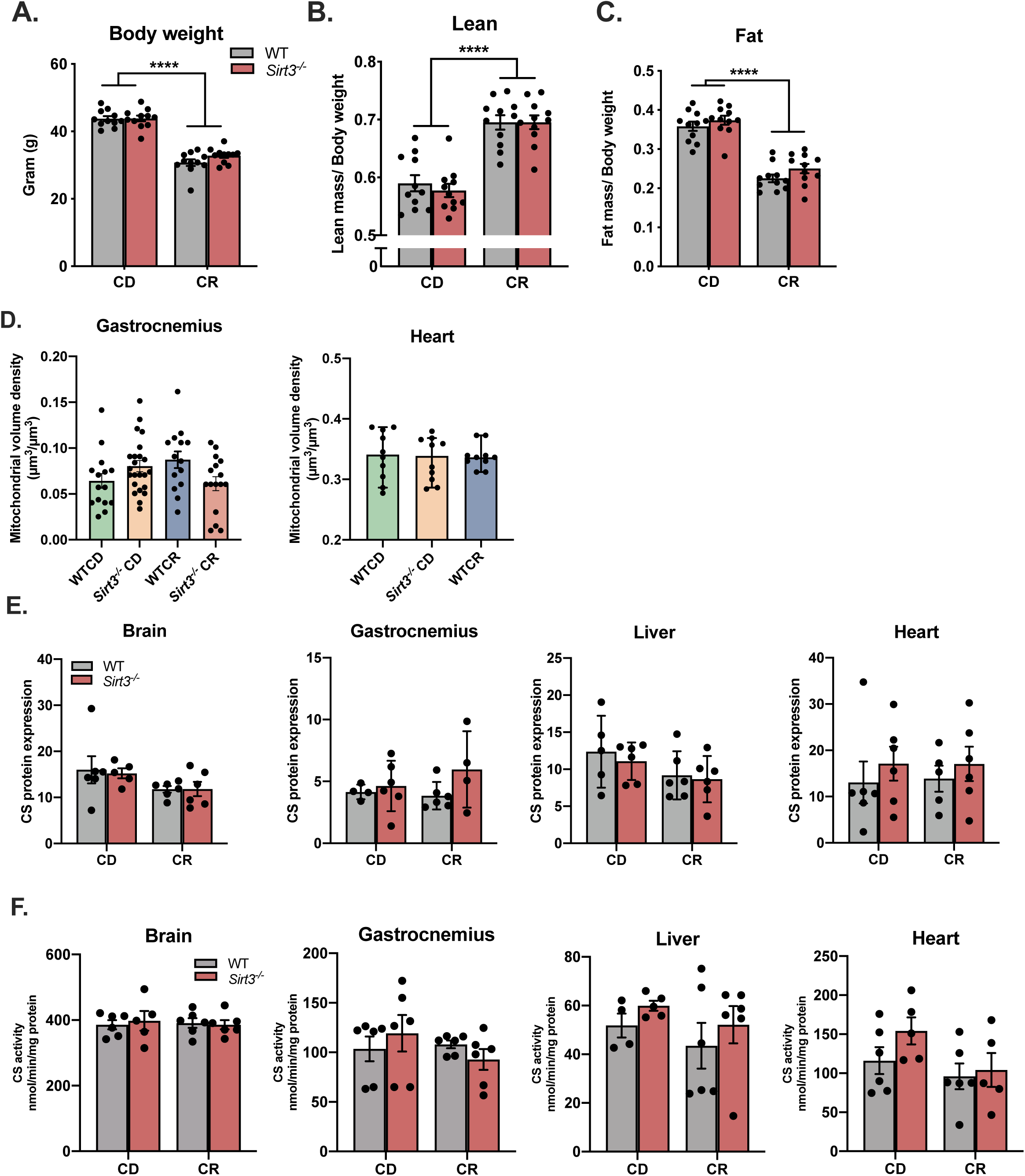
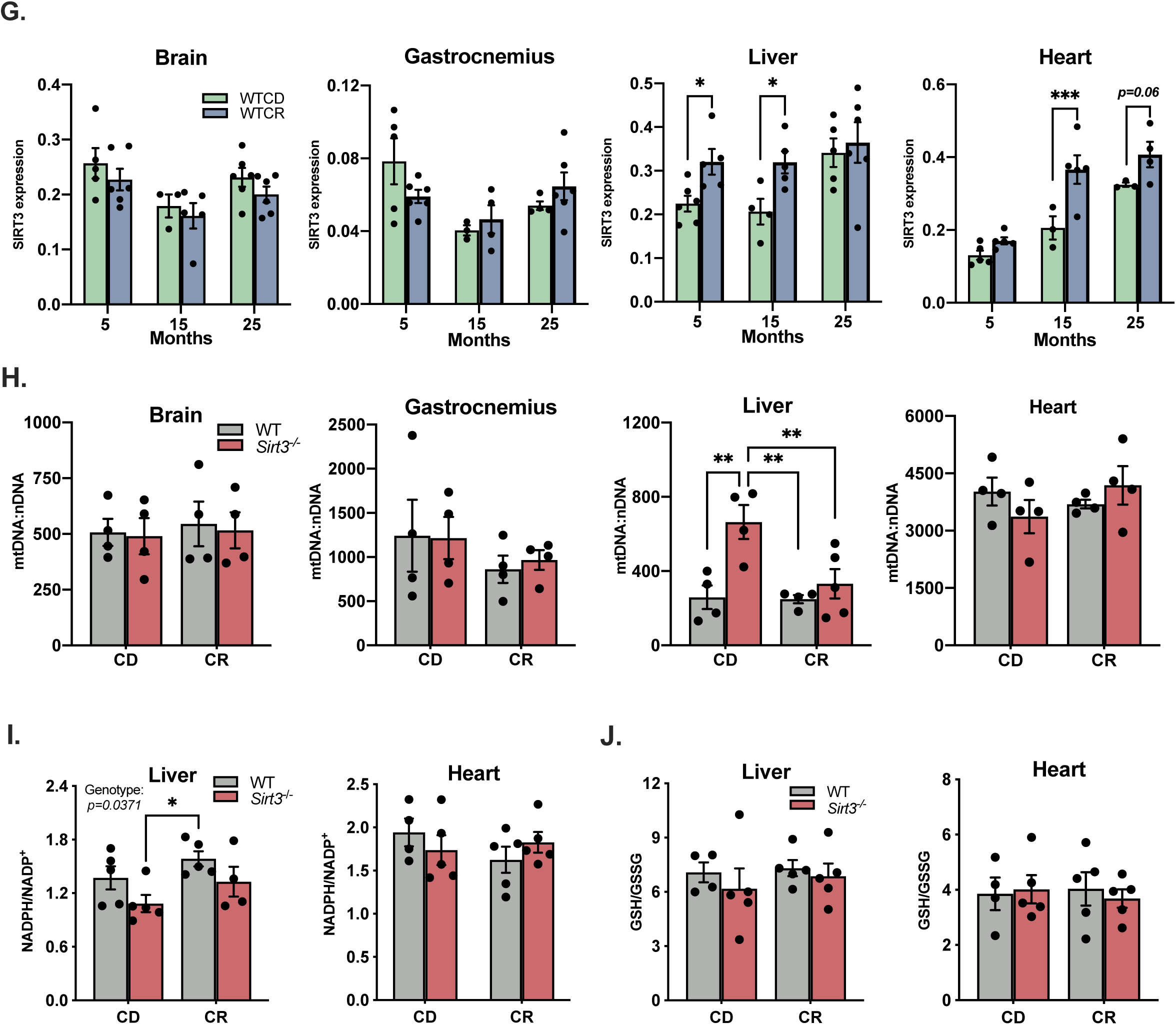
A-C) Bodyweight and body composition for 25-month-old treatment groups, n=11 per group. D) Gastrocnemius and heart mitochondrial volume density from 25-month-old mice. E-F) Citrate synthase expression and activity for 25-month-old treatment groups, n=5-6 per group. Citrate synthase expression was measured using automated capillary electrophoresis-based immunodetection system (WES, ProteinSimple). G) SIRT3 protein expression for 5-, 15-, and 25-month-old WTCD and WTCR groups, n=4-6 per group. SIRT3 expression was measured using automated capillary electrophoresis-based immunodetection system (WES, ProteinSimple). H) Ratio of mitochondrial DNA to nuclear DNA (mDNA:nDNA) for 25-month-old treatment groups, n=4 per group. I-J) NADPH/NADP^+^ and GSH/GSSG in liver and heart from 25-month-old treatment groups measured by LC-MS. NADP(H) and GSH(GSSG) standard curves were used for ratio calculation. n= 4-5 per group. Data were analyzed by two-way ANOVA followed by multiple comparisons test. *p* value reported for each comparison is corrected by Tukey’s test. Results are plotted as mean ± SEM. *: p≤0.05; **: p≤0.01; ***: p≤0.001.

**Fig S2.**
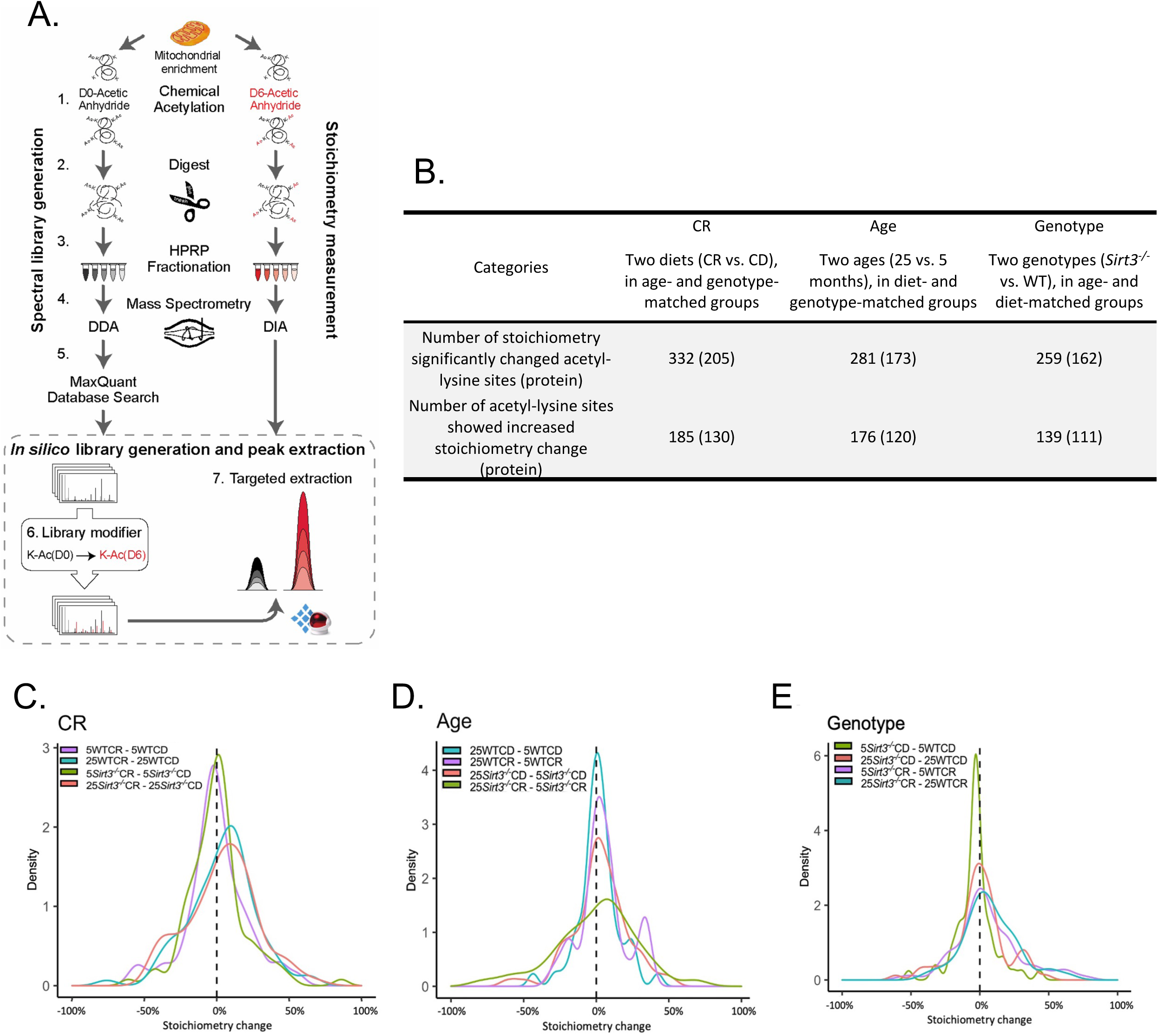
A) Workflow of DIA-dependent acetylation stoichiometry quantification. B) Number of stoichiometry significantly changed (p≤0.05) acetyl-lysine sites and number of acetyl-lysine sites showed increased (>0%) stoichiometry change per factor. Number of proteins are indicated in parentheses. C-E) Distribution of CR-, age-, and genotype-dependent acetylation stoichiometry changes. CR-induced acetylation stoichiometry change is obtained from the stoichiometry difference between CR and CD, in age- and genotype- matched animals. Age-induced acetylation stoichiometry change is obtained from the stoichiometry difference between 25 months and 5 months, in diet- and genotype-matched animals. Genotype-induced acetylation stoichiometry change is obtained from the stoichiometry difference between Sirt3^-/-^ and WT, in diet- and age-matched animals. Only significantly changed acetylation sites (p<0.05) between two groups comparison are included in the density plots. n=4 per group.

**Fig S3.**
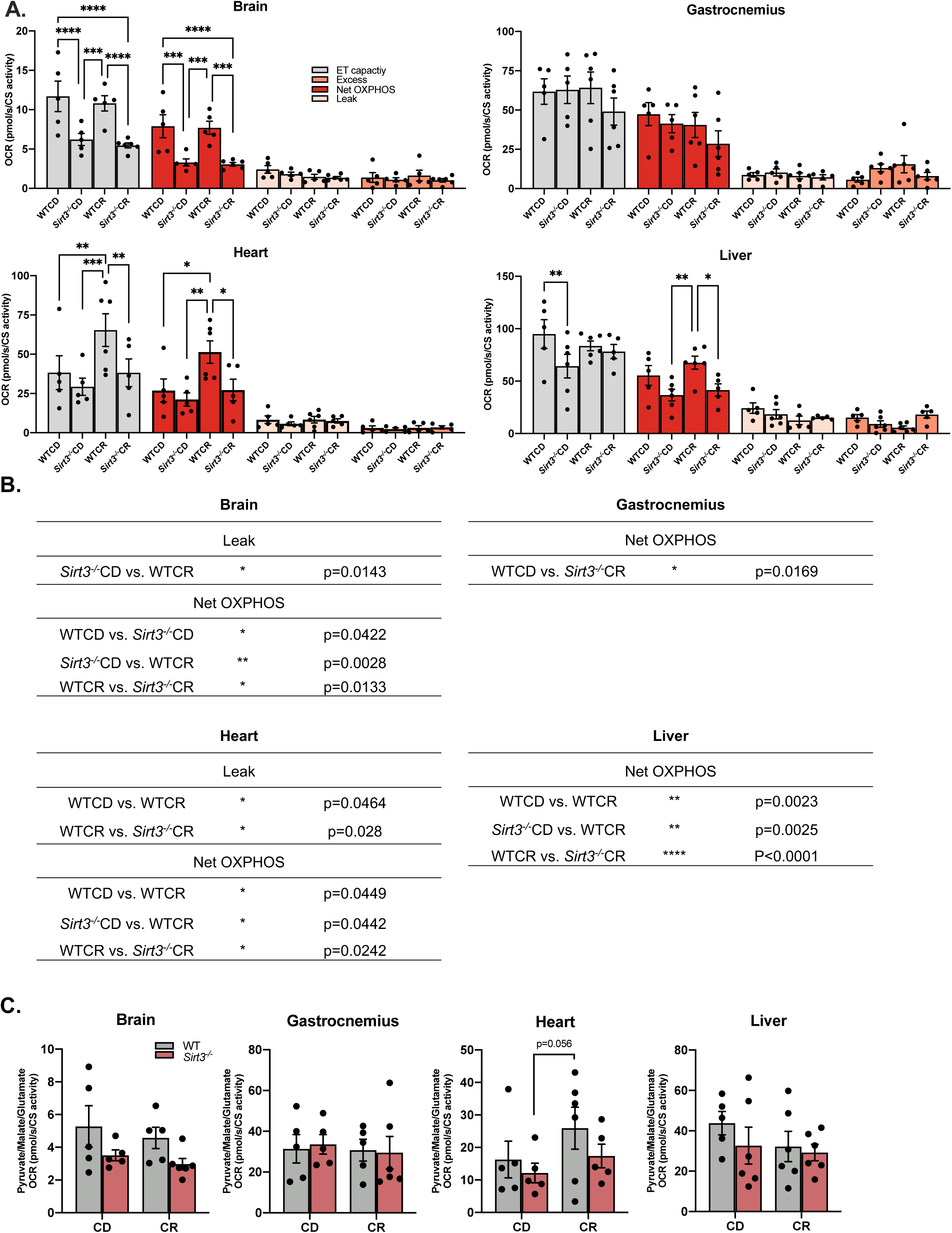
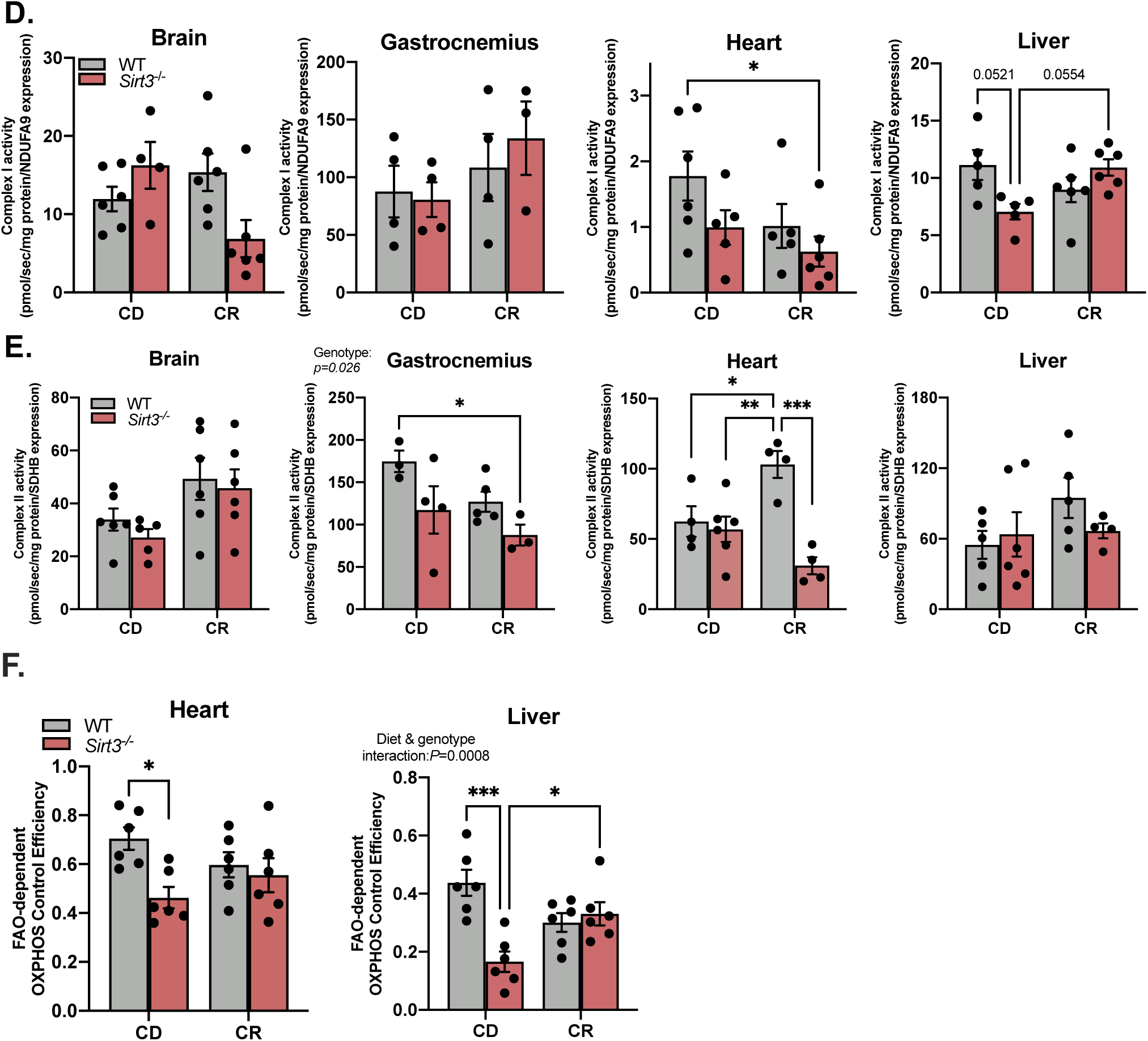
A) Mitochondrial electron transfer (ET) capacity, net OXPHOS, leak and excess respiration of permeabilized brain, gastrocnemius, heart and liver from 25-month-old WTCD, *Sirt3^-/-^*CD, WTCR, *Sirt3^-/-^* CR mice, n= 5-6 per group. Each of these respiration parameter was assessed by oxygen consumption rate (OCR). Plotted data were corrected to citrate synthase (CS) activity. These respiration parameters were assessed by oxygen consumption rate (OCR). Electron transfer capacity was assayed using pyruvate/glutamate/malate/succinate as substrates upon mitochondrial uncoupler FCCP addition. Leak respiration was assayed using pyruvate/glutamate/malate in the absence of ADP. Coupled respiration was assayed using pyruvate/glutamate/malate/succinate in the presence of ADP. Net OXPHOS respiration was calculated by subtracting leak respiration from coupled respiration. Excess respiration is calculated by subtracting coupled respiration from electron transfer capacity. B) Statistics table of Fig. 3B. C) NADH-linked coupled respiration was assessed using pyruvate/glutamate/malate as substrates in the presence of ADP in brain, gastrocnemius, heart and liver from 25-month-oldtreatment groups. D-E) Complex I, II enzymatic activity of brain, gastrocnemius, heart and liver from 25-month-old treatment groups. Complex activity was normalized to total protein and NDUFA9 (Complex I) or SDHB (Complex II) protein expression. F) FAO-dependent net OXPHOS control efficiency was obtained by (coupled FAO respiration – leak FAO respiration) /(coupled FAO respiration), where coupled FAO respiration was assessed using palmitoylcarnitine/malate as substrates in the presence of ADP and leak FAO respiration was assessed using palmitoylcarnitine/malate as substrates in the absence of ADP. Data were analyzed by two-way ANOVA followed by multiple t-tests. *p* value reported for each comparison is corrected by Tukey’s test. Results plotted as mean ± SEM. *: p≤0.05; **: p≤0.01; ***: p≤0.001; ****: p≤0.0001. Significant (p≤0.05) diet effect, genotype effect and/or diet and genotype interaction for each experiment are indicated in figures.

**Fig S4.**
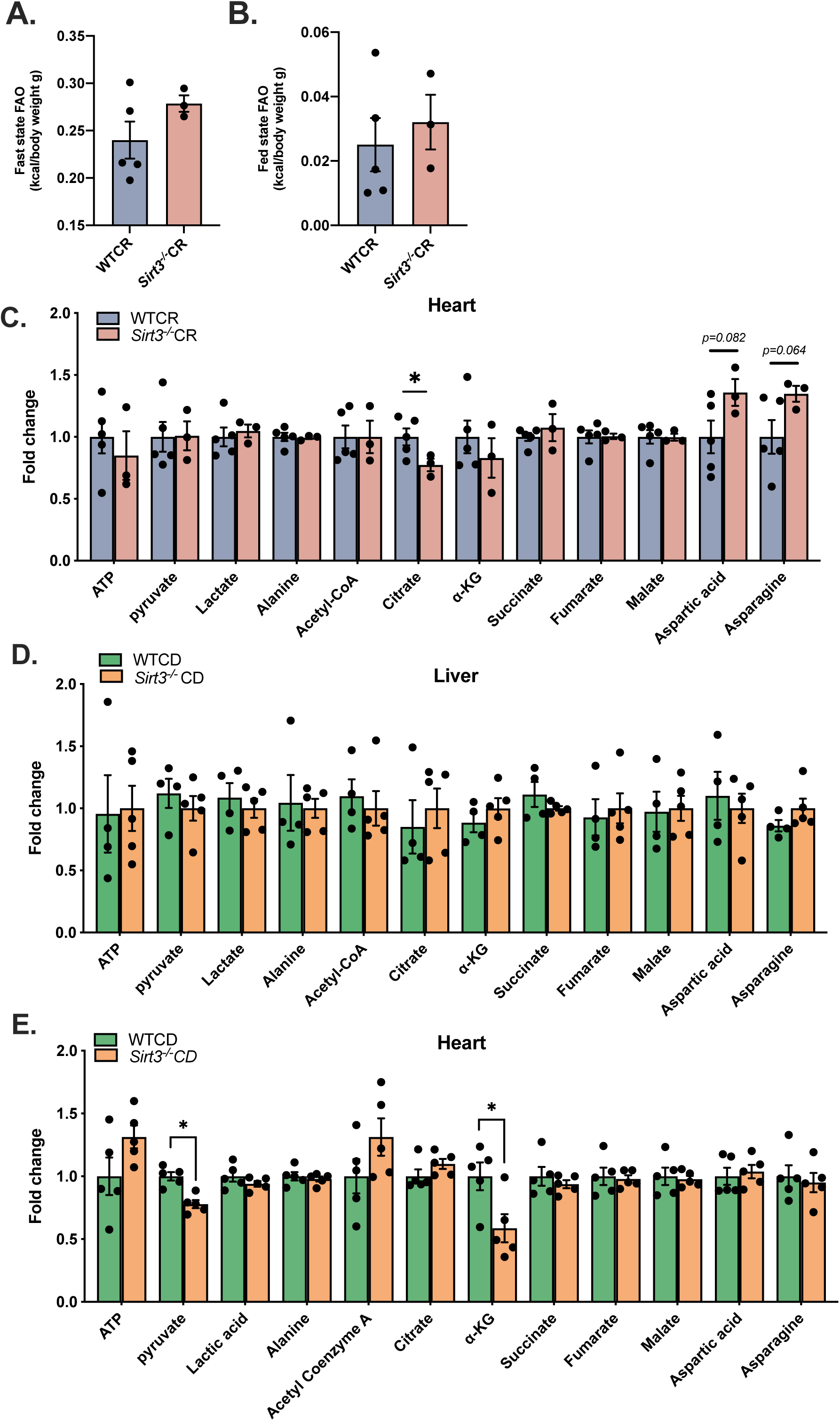
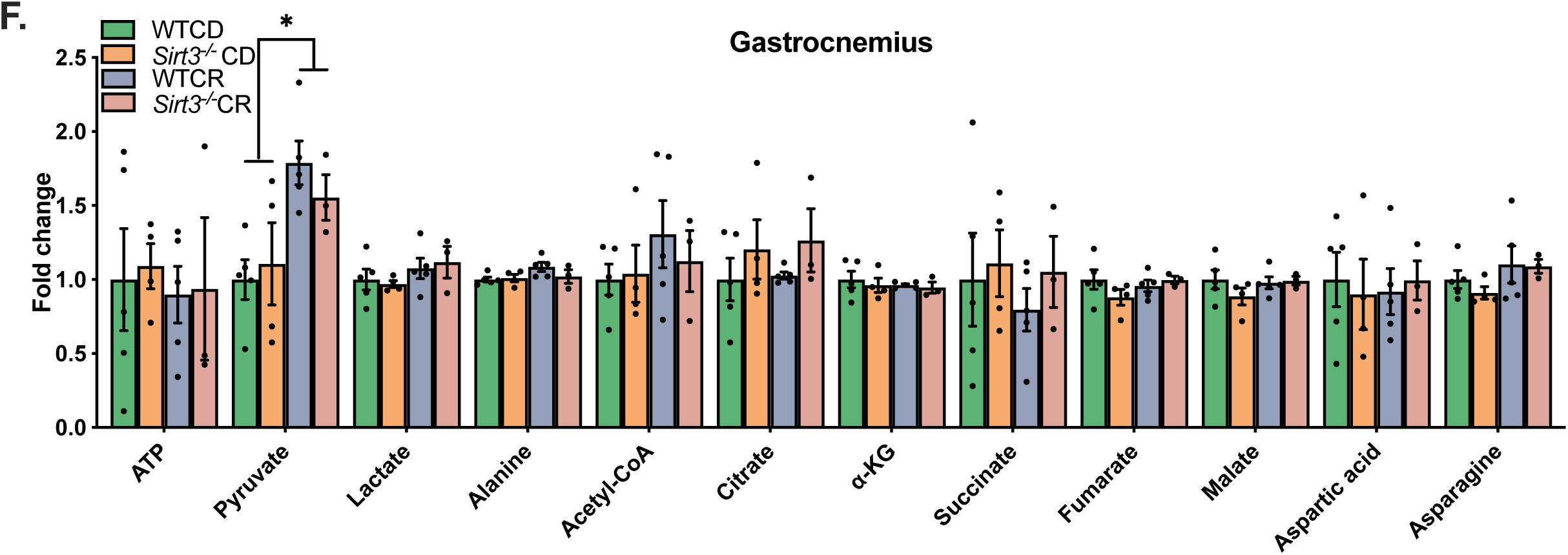
A-B) FAO during fast and fed states for mice shown in Fig.4F. Data reported here are corrected to body weight of individual animals. C-F) Fold change of major TCA metabolites in 25-month-old mice after a 6-hour refeeding, n=3-5. Fig. S4A-E were analyzed by Welch t-test. Fig. S4F was analyzed by One-way ANOVA. Results plotted as mean ± SEM. *: p≤0.05.

**Table S1.**
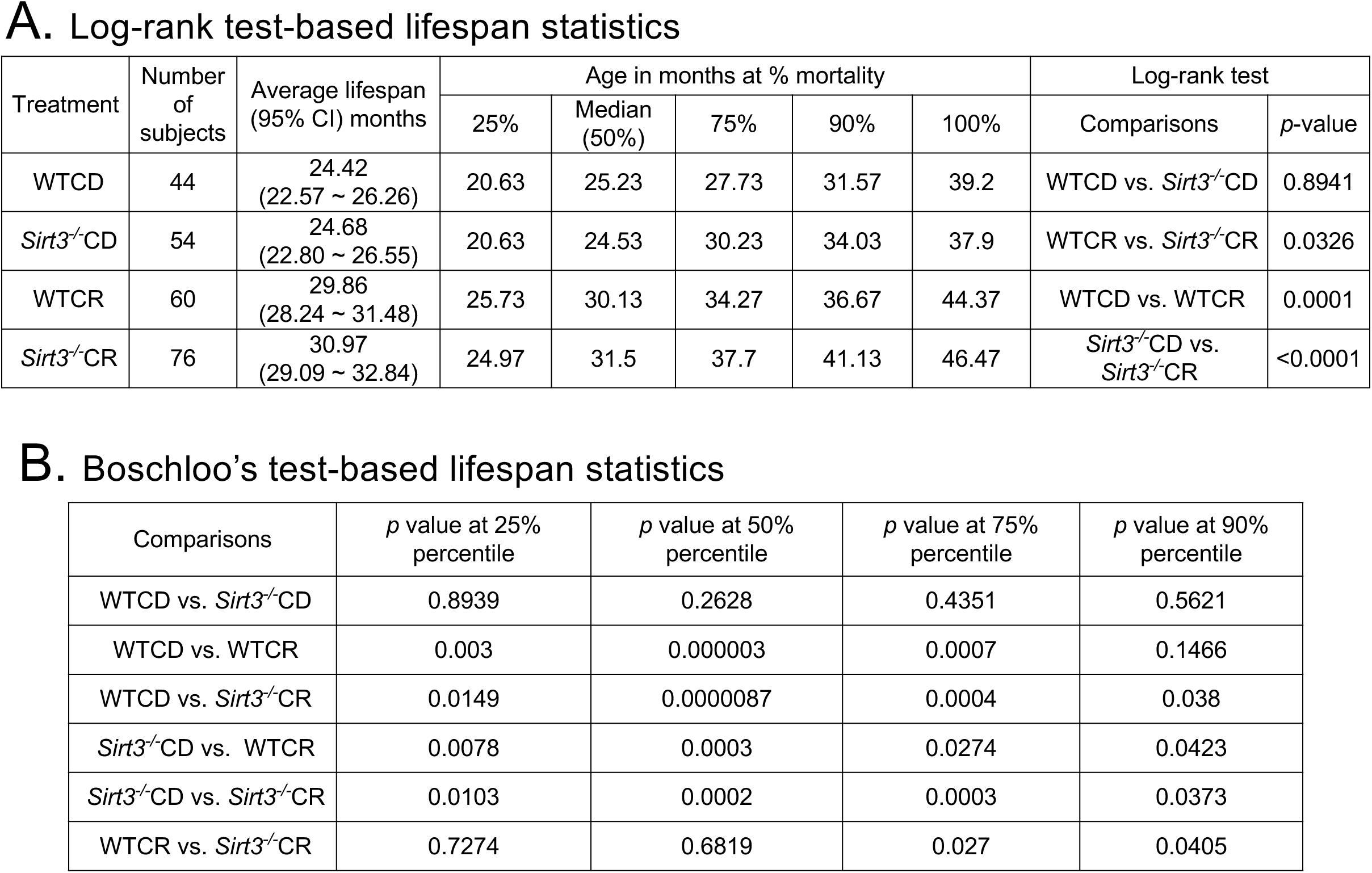
A) Log-rank test-based lifespan statistics of Kaplan-Meier survival curves shown in Fig.1B. B) Boschloo’s test (Wang-Allison)-based lifespan statistics of Kaplan-Meier survival curves shown in Fig.1B. Both Log-rank test and Boschloo’s test were calculated using Online Application for Survival Analysis 2 (OASIS 2) (Han *et al*., 2016).

## REFERENCES

Acosta-Rodríguez, V. A. et al. (2017) ‘Mice under Caloric Restriction Self-Impose a Temporal Restriction of Food Intake as Revealed by an Automated Feeder System’, Cell Metabolism. Cell Press, 26(1), pp. 267–277.e2. doi: 10.1016/j.cmet.2017.06.007.

Agrawal, S. et al. (2019) ‘EL-MAVEN: A fast, robust, and user-friendly mass spectrometry data processing engine for metabolomics’, in *Methods in Molecular Biology*. Methods Mol Biol, pp. 301–321. doi: 10.1007/978-1-4939-9236-2_19.

Ahn, B. H. et al. (2008) ‘A role for the mitochondrial deacetylase Sirt3 in regulating energy homeostasis’, Proceedings of the National Academy of Sciences of the United States of America. National Academy of Sciences, 105(38), pp. 14447–14452. doi: 10.1073/pnas.0803790105.

Ansari, A. et al. (2017) ‘Function of the SIRT3 mitochondrial deacetylase in cellular physiology, cancer, and neurodegenerative disease’, Aging Cell. Blackwell Publishing Ltd, pp. 4–16. doi: 10.1111/acel.12538.

Baeza, J. et al. (2020) ‘Revealing Dynamic Protein Acetylation across Subcellular Compartments’, Journal of Proteome Research. American Chemical Society, 19(6), pp. 2404–2418. doi: 10.1021/acs.jproteome.0c00088.

Baeza, J., Smallegan, M. J. and Denu, J. M. (2016) ‘Mechanisms and Dynamics of Protein Acetylation in Mitochondria’, Trends in Biochemical Sciences. Elsevier Ltd, 41(3), pp. 231–244. doi: 10.1016/j.tibs.2015.12.006.

Balasubramanian, P., Howell, P. R. and Anderson, R. M. (2017) ‘Aging and Caloric Restriction Research: A Biological Perspective With Translational Potential’, EBioMedicine. Elsevier B.V., pp. 37–44. doi: 10.1016/j.ebiom.2017.06.015.

Bao, J. et al. (2010) ‘SIRT3 is regulated by nutrient excess and modulates hepatic susceptibility to lipotoxicity’, Free Radical Biology and Medicine. Pergamon, 49(7), pp. 1230–1237. doi: 10.1016/J.FREERADBIOMED.2010.07.009.

Bartke, A. and Westbrook, R. (2012) ‘Metabolic characteristics of long-lived mice’, Frontiers in Genetics. Frontiers Media SA, pp. 1–6. doi: 10.3389/fgene.2012.00288.

Battaile, K. P. et al. (2004) ‘Structures of Isobutyryl-CoA Dehydrogenase and Enzyme-Product Complex: COMPARISON WITH ISOVALERYL- AND SHORT-CHAIN ACYL-COA DEHYDROGENASES’, Journal of Biological Chemistry. Elsevier, 279(16), pp. 16526–16534. doi: 10.1074/JBC.M400034200.

Bause, A. S. and Haigis, M. C. (2013) ‘SIRT3 regulation of mitochondrial oxidative stress’, Experimental Gerontology, pp. 634–639. doi: 10.1016/j.exger.2012.08.007.

Benigni, A. et al. (2019) ‘Sirt3 Deficiency Shortens Life Span and Impairs Cardiac Mitochondrial Function Rescued by Opa1 Gene Transfer’, Antioxidants and Redox Signaling. Mary Ann Liebert Inc., 31(17), pp. 1255–1271. doi: 10.1089/ars.2018.7703.

Biesiadecki, B. J. et al. (1999) ‘A Gravimetric Method for the Measurement of Total Spontaneous Activity in Rats’, Proceedings of the Society for Experimental Biology and Medicine. Proc Soc Exp Biol Med, 222(1), pp. 65–69. doi: 10.1111/j.1525-1373.1999.09996.x.

Bruss, M. D. et al. (2010) ‘Calorie restriction increases fatty acid synthesis and whole body fat oxidation rates’, American Journal of Physiology-Endocrinology and Metabolism. American Physiological Society Bethesda, MD, 298(1), pp. E108–E116. doi: 10.1152/ajpendo.00524.2009.

Calvo, S. E., Clauser, K. R. and Mootha, V. K. (2016) ‘MitoCarta2.0: An updated inventory of mammalian mitochondrial proteins’, Nucleic Acids Research, 44(D1), pp. D1251–D1257. doi: 10.1093/nar/gkv1003.

Cimen, H. et al. (2010) ‘Regulation of succinate dehydrogenase activity by SIRT3 in mammalian mitochondria’, Biochemistry. NIH Public Access, 49(2), pp. 304–311. doi: 10.1021/bi901627u.

Dennis, G. et al. (2003) ‘DAVID: Database for Annotation, Visualization, and Integrated Discovery.’, Genome biology. BioMed Central, 4(5), pp. 1–11. doi: 10.1186/gb-2003-4-9-r60.

Dittenhafer-Reed, K. E. et al. (2015) ‘SIRT3 mediates multi-tissue coupling for metabolic fuel switching’, Cell Metabolism. Cell Press, 21(4), pp. 637–646. doi: 10.1016/j.cmet.2015.03.007.

Doerrier, C. et al. (2018) ‘High-resolution fluorespirometry and oxphos protocols for human cells, permeabilized fibers from small biopsies of muscle, and isolated mitochondria’, in *Methods in Molecular Biology*. Methods Mol Biol, pp. 31–70. doi: 10.1007/978-1-4939-7831-1_3.

Finley, L. W. S. et al. (2011) ‘Succinate dehydrogenase is a direct target of sirtuin 3 deacetylase activity’, PLoS ONE. Edited by P. Cobine. Public Library of Science, 6(8), p. e23295. doi: 10.1371/journal.pone.0023295.

Fontana, L. and Partridge, L. (2015) ‘Promoting health and longevity through diet: From model organisms to humans’, Cell, pp. 106–118. doi: 10.1016/j.cell.2015.02.020.

Freeman, H. C. et al. (2006) ‘Deletion of nicotinamide nucleotide transhydrogenase: A new quantitive trait locus accounting for glucose intolerance in C57BL/6J mice’, Diabetes. American Diabetes Association, 55(7), pp. 2153–2156. doi: 10.2337/db06-0358.

Fukushima, A. and Lopaschuk, G. D. (2016) ‘Acetylation control of cardiac fatty acid β-oxidation and energy metabolism in obesity, diabetes, and heart failure’, Biochimica et Biophysica Acta - Molecular Basis of Disease. Elsevier B.V., pp. 2211–2220. doi: 10.1016/j.bbadis.2016.07.020.

Gnaiger - MitoEagle Task, E. (2020) ‘Mitochondrial physiology Extended resource of Mitochondrial respiratory states and rates’, Bioenerg Commun, 2020(1), pp. 1–1. doi: 10.26124/bec:2020-0001.v1.

Goetzman, E. S. (2011) ‘Modeling Disorders of Fatty Acid Metabolism in the Mouse’, in *Progress in Molecular Biology and Translational Science*. Elsevier B.V., pp. 389–417. doi: 10.1016/B978-0-12-384878-9.00010-8.

Hallows, W. C. et al. (2011) ‘Sirt3 Promotes the Urea Cycle and Fatty Acid Oxidation during Dietary Restriction’, Molecular Cell. Elsevier Inc., 41(2), pp. 139–149. doi: 10.1016/j.molcel.2011.01.002.

Han, S. K. et al. (2016) ‘OASIS 2: online application for survival analysis 2 with features for the analysis of maximal lifespan and healthspan in aging research’, Oncotarget. Impact Journals, 7(35), pp. 56147–56152. doi: 10.18632/ONCOTARGET.11269.

Haws, S. A. et al. (2020) ‘Methyl-Metabolite Depletion Elicits Adaptive Responses to Support Heterochromatin Stability and Epigenetic Persistence’, Molecular Cell, 78(2), pp. 210–223.e8. doi: 10.1016/j.molcel.2020.03.004.

Hepple, R. T. et al. (2005) ‘Long-term caloric restriction abrogates the age-related decline in skeletal muscle aerobic function’, The FASEB Journal. John Wiley & Sons, Ltd, 19(10), pp. 1320–1322. doi: 10.1096/fj.04-3535fje.

Hirschey, M. D. et al. (2010) ‘SIRT3 regulates mitochondrial fatty-acid oxidation by reversible enzyme deacetylation’, Nature, 464(7285), pp. 121–125. doi: 10.1038/nature08778.

Hofer, T. et al. (2006) ‘A method to determine RNA and DNA oxidation simultaneously by HPLC-ECD: Greater RNA than DNA oxidation in rat liver after doxorubicin administration’, Biological Chemistry. Biol Chem, 387(1), pp. 103–111. doi: 10.1515/BC.2006.014.

Horton, J. L. et al. (2016) ‘Mitochondrial protein hyperacetylation in the failing heart’, JCI Insight. American Society for Clinical Investigation, 1(2). doi: 10.1172/jci.insight.84897.

Jang, J. Y. et al. (2018) ‘The role of mitochondria in aging’, Journal of Clinical Investigation. American Society for Clinical Investigation, pp. 3662–3670. doi: 10.1172/JCI120842.

Jing, E. et al. (2013) ‘Sirt3 regulates metabolic flexibility of skeletal muscle through reversible enzymatic deacetylation’, Diabetes, 62(10), pp. 3404–3417. doi: 10.2337/db12-1650.

Joseph, A. M. et al. (2012) ‘The impact of aging on mitochondrial function and biogenesis pathways in skeletal muscle of sedentary high- and low-functioning elderly individuals’, Aging Cell. NIH Public Access, 11(5), pp. 801–809. doi: 10.1111/j.1474-9726.2012.00844.x.

Kim, H. S. et al. (2010) ‘SIRT3 Is a Mitochondria-Localized Tumor Suppressor Required for Maintenance of Mitochondrial Integrity and Metabolism during Stress’, Cancer Cell. Elsevier Ltd, 17(1), pp. 41–52. doi: 10.1016/j.ccr.2009.11.023.

Kincaid, B. and Bossy-Wetzel, E. (2013) ‘Forever young: SIRT3 a shield against mitochondrial meltdown, aging, and neurodegeneration’, Frontiers in Aging Neuroscience. Frontiers Media SA, p. 48. doi: 10.3389/fnagi.2013.00048.

Kľučková, K. et al. (2020) ‘Succinate dehydrogenase deficiency in a chromaffin cell model retains metabolic fitness through the maintenance of mitochondrial NADH oxidoreductase function’, The FASEB Journal, 34(1), pp. 303–315. doi: 10.1096/fj.201901456R.

Korasick, D. A. et al. (2018) ‘NAD+ promotes assembly of the active tetramer of aldehyde dehydrogenase 7A1’, FEBS Letters. John Wiley & Sons, Ltd, 592(19), pp. 3229–3238. doi: 10.1002/1873-3468.13238.

Kurimoto, K. et al. (2001) ‘Crystal Structure of Human AUH Protein, a Single-Stranded RNA Binding Homolog of Enoyl-CoA Hydratase’, Structure. Cell Press, 9(12), pp. 1253– 1263. doi: 10.1016/S0969-2126(01)00686-4.

Kuznetsov, A. V et al. (2008) ‘Analysis of mitochondrial function in situ in permeabilized muscle fibers, tissues and cells’, Nature Protocols. Nature Publishing Group, 3(6), pp. 965–976. doi: 10.1038/nprot.2008.61.

Lantier, L. et al. (2015) ‘SIRT3 is crucial for maintaining skeletal muscle insulin action and protects against severe insulin resistance in high-fat-fed mice’, Diabetes. American Diabetes Association Inc., 64(9), pp. 3081–3092. doi: 10.2337/db14-1810.

Lanza, I. R. et al. (2012) ‘Chronic caloric restriction preserves mitochondrial function in senescence without increasing mitochondrial biogenesis’, Cell Metabolism. NIH Public Access, 16(6), pp. 777–788. doi: 10.1016/j.cmet.2012.11.003.

Latorre-Muro, P. et al. (2018) ‘Dynamic Acetylation of Phosphoenolpyruvate Carboxykinase Toggles Enzyme Activity between Gluconeogenic and Anaplerotic Reactions’, Molecular Cell. Elsevier, 71(5), pp. 718–732.e9. doi: 10.1016/j.molcel.2018.07.031.

Leibel R, Rosenbaum M, H. J. (2020) ‘Flux through NNT-linked mitochondrial redox buffering circuits generates counterbalance changes in energy expenditure’, JBC, 332, pp. 621–8.

Liu, J. et al. (2017) ‘Sirt3 protects hepatocytes from oxidative injury by enhancing ros scavenging and mitochondrial integrity’, Cell Death and Disease. Nature Publishing Group, 8(10), pp. e3158–e3158. doi: 10.1038/cddis.2017.564.

Liu, Z. et al. (2014) ‘CPLM: A database of protein lysine modifications’, Nucleic Acids Research. Nucleic Acids Res, 42(D1). doi: 10.1093/nar/gkt1093.

Luo, M. et al. (2015) ‘Diethylaminobenzaldehyde is a covalent, irreversible inactivator of ALDH7A1’, ACS Chemical Biology. American Chemical Society, 10(3), pp. 693–697. doi: 10.1021/CB500977Q/SUPPL_FILE/CB500977Q_SI_001.PDF.

Manini, T. M. (2010) ‘Energy Expenditure and Aging’, Ageing research reviews. NIH Public Access, 9(1), p. 1. doi: 10.1016/J.ARR.2009.08.002.

Mattison, J. A. et al. (2017) ‘Caloric restriction improves health and survival of rhesus monkeys’, Nature Communications, 8. doi: 10.1038/ncomms14063.

Mattson, M. P. et al. (2014) ‘Meal frequency and timing in health and disease’, Proceedings of the National Academy of Sciences of the United States of America. Proc Natl Acad Sci U S A, 111(47), pp. 16647–16653. doi: 10.1073/pnas.1413965111.

McDonnell, E. et al. (2015) ‘SIRT3 regulates progression and development of diseases of aging’, Trends in Endocrinology and Metabolism, pp. 486–492. doi: 10.1016/j.tem.2015.06.001.

Merry, B. J. (2004) ‘Oxidative stress and mitochondrial function with aging - The effects of calorie restriction’, Aging Cell. John Wiley & Sons, Ltd, pp. 7–12. doi: 10.1046/j.1474-9728.2003.00074.x.

Mezhnina, V. et al. (2020) ‘CR reprograms acetyl-CoA metabolism and induces long-chain acyl-CoA dehydrogenase and CrAT expression’, Aging Cell. Blackwell Publishing Ltd, 19(11). doi: 10.1111/acel.13266.

Mihaylova, M. M. et al. (2018) ‘Fasting Activates Fatty Acid Oxidation to Enhance Intestinal Stem Cell Function during Homeostasis and Aging’, Cell Stem Cell. Cell Stem Cell, 22(5), pp. 769–778.e4. doi: 10.1016/j.stem.2018.04.001.

Mitchell, S. J. et al. (2019) ‘Daily Fasting Improves Health and Survival in Male Mice Independent of Diet Composition and Calories’, Cell Metabolism, 29, pp. 221–228. doi: 10.1016/j.cmet.2018.08.011.

Ono, K. et al. (2019) ‘SIRT3 Regulation of Mitochondrial Quality Control in Neurodegenerative Diseases’, Front. Aging Neurosci, 11, p. 313. doi: 10.3389/fnagi.2019.00313.

Pak, H. H. et al. (2021) ‘Distinct roles of fasting and calories in the metabolic, molecular, and geroprotective effects of a calorie restricted diet’, Nature Metabolism. doi: NATMETAB-A21014120B.

Palacios, O. M. et al. (2009) ‘Diet and exercise signals regulate SIRT3 and activate AMPK and PGC-1alpha in skeletal muscle.’, Aging. Impact Journals, LLC, 1(9), pp. 771–783. doi: 10.18632/aging.100075.

Parodi-Rullán, R. M., Chapa-Dubocq, X. R. and Javadov, S. (2018) ‘Acetylation of Mitochondrial Proteins in the Heart: The Role of SIRT3’, Frontiers in Physiology. Frontiers, 9, p. 1094. doi: 10.3389/fphys.2018.01094.

Patil, Y. N. et al. (2015) ‘Cellular and molecular remodeling of inguinal adipose tissue mitochondria by dietary methionine restriction’, Journal of Nutritional Biochemistry. J Nutr Biochem, 26(11), pp. 1235–1247. doi: 10.1016/j.jnutbio.2015.05.016.

Pougovkina, O. et al. (2014) ‘Mitochondrial protein acetylation is driven by acetyl-CoA from fatty acid oxidation’, Human Molecular Genetics. Oxford University Press, 23(13), pp. 3513–3522. doi: 10.1093/hmg/ddu059.

Pugh, T. D., Klopp, R. G. and Weindruch, R. (1999) ‘Controlling caloric consumption: Protocols for rodents and rhesus monkeys’, Neurobiology of Aging, 20(2), pp. 157–165. doi: 10.1016/S0197-4580(99)00043-3.

Qiu, X. et al. (2010) ‘Calorie restriction reduces oxidative stress by SIRT3-mediated SOD2 activation’, Cell Metabolism, 12(6), pp. 662–667. doi: 10.1016/j.cmet.2010.11.015.

Richardson, N. E. et al. (2021) ‘Lifelong restriction of dietary branched-chain amino acids has sex-specific benefits for frailty and life span in mice’, Nature Aging 2020 1:1. Nature Publishing Group, 1(1), pp. 73–86. doi: 10.1038/s43587-020-00006-2.

Ronchi, J. A. et al. (2016) ‘The Contribution of Nicotinamide Nucleotide Transhydrogenase to Peroxide Detoxification Is Dependent on the Respiratory State and Counterbalanced by Other Sources of NADPH in Liver Mitochondria’, Journal of Biological Chemistry, 291(38), pp. 20173–20187. doi: 10.1074/jbc.M116.730473.

Scariot, P. P. M. et al. (2016) ‘Continuous aerobic training in individualized intensity avoids spontaneous physical activity decline and improves MCT1 expression in oxidative muscle of swimming rats’, *Frontiers in Physiology*. Front Physiol, 7(APR). doi: 10.3389/fphys.2016.00132.

Scariot, P. P. M. et al. (2019) ‘Housing conditions modulate spontaneous physical activity, feeding behavior, aerobic running capacity and adiposity in C57BL/6J mice’, Hormones and Behavior. Elsevier, 115(July), p. 104556. doi: 10.1016/j.yhbeh.2019.07.004.

Schmiedl, A. et al. (1990) ‘The surface to volume ratio of mitochondria, a suitable parameter for evaluating mitochondrial swelling - Correlations during the course of myocardial global ischaemia’, *Virchows Archiv A Pathological Anatomy and Histopathology*. Virchows Arch A Pathol Anat Histopathol, 416(4), pp. 305–315. doi: 10.1007/BF01605291.

Schwer, B. et al. (2009) ‘Calorie restriction alters mitochondrial protein acetylation’, Aging Cell. NIH Public Access, pp. 604–606. doi: 10.1111/j.1474-9726.2009.00503.x.

Seiler, S. E. et al. (2015) ‘Carnitine Acetyltransferase Mitigates Metabolic Inertia and Muscle Fatigue during Exercise’, Cell Metabolism. Cell Press, 22(1), pp. 65–76. doi: 10.1016/j.cmet.2015.06.003.

Someya, S. et al. (2010) ‘Sirt3 mediates reduction of oxidative damage and prevention of age-related hearing loss under Caloric Restriction’, Cell. Elsevier Inc., 143(5), pp. 802–812. doi: 10.1016/j.cell.2010.10.002.

Srivastava, S. (2017) ‘The mitochondrial basis of aging and age-related disorders’, Genes. MDPI AG, p. 398. doi: 10.3390/genes8120398.

Still, A. J. et al. (2013) ‘Quantification of Mitochondrial Acetylation Dynamics Highlights Prominent Sites of Metabolic Regulation’, The Journal of Biological Chemistry. American Society for Biochemistry and Molecular Biology, 288(36), p. 26209. doi: 10.1074/JBC.M113.483396.

Talal, S. et al. (2021) ‘High carbohydrate diet ingestion increases post-meal lipid synthesis and drives respiratory exchange ratios above 1’, Journal of Experimental Biology. The Company of Biologists, 224(4). doi: 10.1242/JEB.240010.

Tsuda, M. et al. (2018) ‘Protein acetylation in skeletal muscle mitochondria is involved in impaired fatty acid oxidation and exercise intolerance in heart failure’, Journal of Cachexia, Sarcopenia and Muscle. Springer Nature, 9(5), pp. 844–859. doi: 10.1002/jcsm.12322.

Ullman-Culleré, M. H. and Foltz, C. J. (1999) Body Condition Scoring: A Rapid and Accurate Method for Assessing Health Status in Mice.

Vassilopoulos, A. et al. (2014) ‘SIRT3 deacetylates ATP synthase F1 complex proteins in response to nutrient-and exercise-induced stress’, Antioxidants and Redox Signaling. Mary Ann Liebert, Inc. 140 Huguenot Street, 3rd Floor New Rochelle, NY 10801 USA, 21(4), pp. 551–564. doi: 10.1089/ars.2013.5420.

Weindruch, R. et al. (1986) ‘The retardation of aging in mice by dietary restriction: Longevity, cancer, immunity and lifetime energy intake’, Journal of Nutrition. Oxford Academic, 116(4), pp. 641–654. doi: 10.1093/jn/116.4.641.

Weindruch, R. H. et al. (1980) ‘Modification of mitochondrial respiration by aging and dietary restriction’, Mechanisms of Ageing and Development. Elsevier, 12(4), pp. 375– 392. doi: 10.1016/0047-6374(80)90070-6.

Weindruch, R. and Sohal, R. S. (1997) ‘Caloric Intake and Aging’, New England Journal of Medicine. NIH Public Access, 337(14), pp. 986–994. doi: 10.1056/nejm199710023371407.

Williams, A. S. et al. (2019) ‘Disruption of Acetyl-Lysine Turnover in Muscle Mitochondria Promotes Insulin Resistance and Redox Stress without Overt Respiratory Dysfunction’, Cell Metabolism, 31(1), pp. 131–147.e11. doi: 10.1016/j.cmet.2019.11.003.

Yu, D. et al. (2021) ‘The adverse metabolic effects of branched-chain amino acids are mediated by isoleucine and valine’, Cell Metabolism. Cell Press, 33(5), pp. 905–922.e6. doi: 10.1016/J.CMET.2021.03.025.

Yu, W., Dittenhafer-Reed, K. E. and Denu, J. M. (2012) ‘SIRT3 protein deacetylates isocitrate dehydrogenase 2 (IDH2) and regulates mitochondrial redox status’, Journal of Biological Chemistry. American Society for Biochemistry and Molecular Biology, 287(17), pp. 14078–14086. doi: 10.1074/jbc.M112.355206.

Zeng, L. et al. (2014) ‘Age-related decrease in the mitochondrial sirtuin deacetylase sirt3 expression associated with ROS accumulation in the auditory cortex of the mimetic aging rat model’, PLoS ONE. Public Library of Science, 9(2). doi: 10.1371/journal.pone.0088019.

Zhang, C. et al. (2013) ‘Structural modulation of gut microbiota in life-long calorie-restricted mice’, Nature Communications. Nature Publishing Group, 4(1), pp. 1–10. doi: 10.1038/ncomms3163.

